# Collective fluctuations underlying nanobody inhibitory activity targeting *B. anthracis* S-layers revealed by multiscale simulations

**DOI:** 10.64898/2026.06.26.734918

**Authors:** Adam J. Cecil, Alexander J. Pak

**Affiliations:** Department of Chemical and Biological Engineering, Colorado School of Mines, Golden, CO, USA; Quantitative Biosciences and Engineering Program, Colorado School of Mines, Golden, CO, USA; Materials Science Program, Colorado School of Mines, Golden, CO, USA

## Abstract

Nanobodies (Nbs) that depolymerize bacterial surface-layers (S-layers) offer a route to antivirulence therapeutics, but their mechanisms have been difficult to infer from binding structures alone. In *B. anthracis,* several Nbs bind to Sap, the S-layer protein that assembles into a paracrystalline lattice surrounding the cell. Despite similar binding poses and sequences, only a subset of these (“inhibitory”) Nbs induce disassembly of the protein lattice, ultimately abrogating pathogenicity. In this study, we leveraged multiscale simulations to test whether representative Nb-induced fluctuations local to the binding site are sufficient to reproduce and explain lattice-scale depolymerization. Using a divide-and-conquer strategy, we first developed a bottom-up coarse-grained (CG) model of the multidomain Sap monomer. We then compared machine learning (ML) and information theoretic approaches to identify Nb-induced collective fluctuations that are predictive and potentially causative for depolymerization. We found that motions encoding Nb rigidification and partial clamping of the binding site instigate early-stage Sap depolymerization when propagated into lattice-scale computational depolymerization assays. Furthermore, the model informed by correlation-grouped ML analysis correctly reproduced both inhibitory and non-inhibitory phenotypes for 10 out of 12 (Nb)-Sap systems. Combined time-resolved defect and strain analyses revealed that inhibitory Nbs cooperatively apply a critical amount of tensile stress that destabilizes Sap-Sap interfaces parallel to the direction of strain and proximal to the Nb binding site, thereby supporting a mechanism in which local, Nb-imposed fluctuations propagate into lattice-scale mechanical instability. More broadly, this work demonstrates how multiscale simulations combined with ML analysis can test whether molecular-scale conformational signatures are sufficient to drive emergent phenotypes in large protein assemblies. In the future, this general approach can be adapted for mechanistic study and subsequent rational design of therapeutics that rely on dynamical interventions of protein virulence factors, such as through rigidification or assembly-disrupting modes of action.

## Introduction

Antibiotic resistance is one of the top global health threats, and its accelerating development threatens the usefulness of existing antibiotics [1]. Over the last century, antibiotics consistently targeting the same bacterial viability pathways have put extreme selective pressure on pathogens, rapidly progressing their antibiotic resistance [2–4]. Rather than targeting viability, a growing body of research is exploring the possibility of targeting virulence factors instead [2, 5–9]. Virulence factors are small molecules or proteins that bacteria rely on to invade, colonize, and harm their host. By targeting virulence, a therapeutic non-lethally disarms the pathogen, reducing (or potentially, eliminating) the selective pressures that lead to antibiotic resistance [9–12]. One such virulence factor is the surface-layer, or S-layer. S-layers are self-assembling paracrystalline protein lattices present on the surface of many bacteria and almost all archaea [13]. Hypothesized to be products of convergent evolution [14], S-layers are highly specific to each organism and are involved in processes including cell stability, colonization, motility, and environmental signaling, many of which play direct roles in virulence and are not always required for cell viability [13–22]. Their high sequence and structure specificity makes S-layers attractive for targeted therapeutic intervention without disturbing surrounding commensals. However, that same specificity means that S-layer-targeted therapeutics will likely require organism-specific molecular designs, making it essential to identify the mechanism of action in each system and determine which features generalize across S-layer families.

*B. anthracis*, the causative agent of anthrax infections, is a gram-positive bacterium whose S-layer provides mechanical stabilization and takes in environmental signals that trigger virulence [20, 23, 24]. During the exponential growth phase, the S-layer is made of monomers called Sap (<u>s</u>urface <u>a</u>rray <u>p</u>rotein) that form elliptical rings spanning roughly 11×8 nm^2^ in the lattice plane and non-covalently self-assemble into a continuous 2D lattice [23]. The assembly domain of each monomer consists of 6 immunoglobulin (Ig)-like domains (termed “D1” through “D6”) connected by flexible linkers [20, 23, 24]. In 2019, Fioravanti and coworkers developed single-domain camelid antibodies, called nanobodies (Nbs), that disassemble (or “depolymerize”) Sap and non-lethally disarm *B. anthracis* infections [24]. These Nbs bind to a region of Sap that does not directly interfere with Sap-Sap lattice contacts, yet their inhibitory effect is expressed through lattice-scale disassembly. In this study, we set out to uncover the conformational and mechanical steps that connect inhibitory Nb binding to Sap depolymerization, such that in the future one might rationally design better Nb therapeutics targeting *B. anthracis* and related pathogens.

The discovered Nbs bind to the interior of the Sap ring, specifically to the hinge region between the first two domains, D1D2 (**Fig 1A**). Upon binding, the Nbs initiate depolymerization of the lattice (if formed) and prevent Sap assembly (if lattice unformed), both of which ultimately disarm the bacteria. However, of the 11 isolated Nbs that bind with high affinity, only 5 cause depolymerization (which we term “inhibitory Nbs”), and those to varying degrees. Inspection of the complementarity determining region 3 (CDR3) loops responsible for binding reveals chemically similar sequences between both inhibitory and non-inhibitory Nbs [25]. Available X-ray structures of two inhibitory Nbs and one non-inhibitory Nb bound to D1D2 reveal near-identical binding poses. Experimental depolymerization assays have successfully quantified how much Sap each Nb depolymerizes [25], but have yet to resolve the causal chain connecting near-identical Nb-Sap binding to large-scale disassembly.

**Figure 1.**
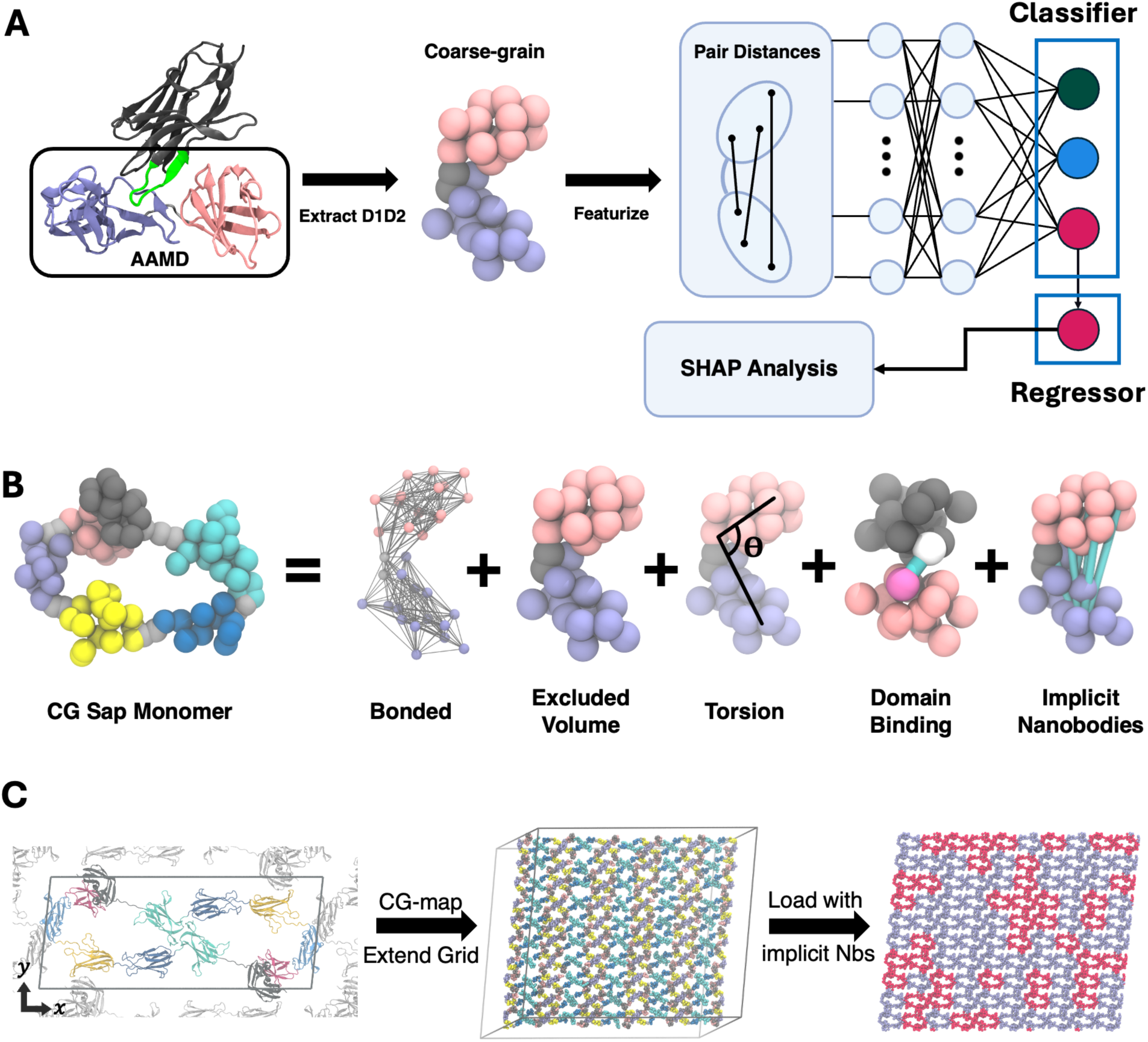
A) MD-ML pipeline to identify the most predictive inter-domain CG pair distances for Nb-induced inhibition. All-atom protein is shown in New Cartoon representation, where pink, ice blue, and black (green) color is D1, D2, and Nb (CDR3 of Nb), respectively. The CG Sap protein is shown as space-filling beads where pink, ice blue, and gray represent D1, D2, and linker beads, respectively. The FFNN outputs 3 class probabilities, where green, blue, and red represent non-bound, non-inhibitory, and inhibitory, respectively. B) Schematic of the CG model formulation, which consists of five terms. Domains are colored as follows: D1 is pink, D2 is ice blue, D3 is yellow, D4 is navy, D5 is cyan, D6 is dark grey, and all linkers are light grey. The white bead in “Domain Binding” represents a VS bonded to the magenta bead via the cyan bond. The cyan bonds in “Implicit Nanobodies” represent the pairwise fluctuations enforced by tabulated bond potentials. C) The lattice unit cell (left) is replicated in *xy* to the supercell or simulation box (center), then loaded with implicit Nbs (right, 40% loading shown here). Ice blue monomers indicate non-bound Sap, and red monomers indicate Nb-bound Sap.

Because the behavior we are interested in is inherently multiscale (spanning Å-scale binding interactions to nm/µm-scale phenotypes), we turn to multiscale modeling where we first aim to probe the atomic interactions that govern Sap’s local conformational behavior with and without Nbs, then propagate those interactions at the scale of assembled Sap lattices. Molecular dynamics (MD) is well-suited to this type of approach. Conventional all-atom MD (AAMD) simulates every atom in a system using classical mechanics and has been used successfully to study the dynamics and self-assembly of peptides and small protein systems [26, 27]. However, even with specialized supercomputers, AAMD is limited to a few hundred nanometers at the microsecond timescale [28]. S-layers like Sap span thousands of nanometers and assemble/depolymerize over minutes to hours [24]. Enhanced-sampling methods raise the probability of sampling rare and slow events to recover free energy profiles and have successfully been used for self-assembly studies [29, 30]. However, while they accelerate sampling of configurational space, the computational cost remains prohibitively high for systems as large as S-layers.

Coarse-grained MD (CGMD) extends both accessible time- and length-scales by grouping atoms together into CG sites that retain the average behavior of their atomic groups and have been applied to large lipid bilayers and other macromolecular assemblies to great effect [31–33]. In general, there are two prevailing CG philosophies: top-down and bottom-up. Top-down methods reproduce macroscopic or thermodynamic properties but are not guaranteed to reproduce microscopic fluctuations. Bottom-up methods explicitly enforce microscopic fluctuations, and only when the correct set of fluctuations is included do emergent macroscopic observables appear. As such, bottom-up CG models act as hypothesis testing frameworks – the model builder hypothesizes which microscopic interactions are sufficient to produce the target macroscopic behavior, and positive results only occur when that behavior emerges in CGMD. Bottom-up CG methods are therefore suitable for mechanism discovery rooted in microscopic fluctuations. However, due to the complexity of large protein assemblies, identifying that sufficient set of microscopic interactions can be highly non-trivial, as the interactions driving emergent behavior may not be obvious, even to domain experts.

In practice, bottom-up CG models are often built in stages. Core interactions that capture established physical behavior are derived using established systematic methods such as multiscale coarse-graining (MS-CG) [34–36] and heterogeneous elastic network models (hENMs) [37], while additional interaction terms are introduced to represent system-specific mechanisms [31, 32, 38–43]. In the case of Sap, a bottom-up CG model should retain the rigidity of individual Ig-like domains, the conformational flexibility of the multidomain protein, and the likely reversible protein-protein interactions responsible for lattice assembly. These features are well motivated by prior studies of proteins and their assemblies [31, 32, 34–37, 39, 41–47]. The greater challenge is determining how Nb binding alters Sap dynamics at the microscopic level in a manner that ultimately triggers lattice disassembly. In prior work, we showed that AAMD simulations of Nb-bound Sap reveal dynamical signatures that are predictive of Sap depolymerization activity [25]. By simulating the Sap binding site (i.e., D1D2) both alone and bound to each of the Nbs, machine learning (ML) models distinguished between the microscopic binding site fluctuations in non-bound, non-inhibitory, and inhibitory-Nb-bound systems. By ranking the input features in terms of their contribution to the ML model, we identified three conformational motions that correlate well with Nb-induced depolymerization. Correlation, however, does not imply a causative mechanism. Therefore, a natural next step is to test whether identified microscopic fluctuations lead to Nb-induced depolymerization in the context of CGMD simulations.

In this study, we tested whether inhibitory-predictive binding site correlations are sufficient to reproduce the phenotypic effects of each Nb at the scale of periodic Sap lattices using CGMD. First, we developed a divide-and-conquer strategy to build a Sap monomer CG model to make parameterization of such a large protein tractable. We then compared three data-driven methods to identify candidate sets of pairwise fluctuation correlations (local to the binding site) that are predicted to distinguish non-bound, non-inhibitory, and inhibitory activity. These selected correlations were converted into tabulated bond potentials and enforced in periodic CG Sap lattices to represent Nb-specific changes to the binding site. Using this framework, we quantified defect formation, defect propagation, lattice strain, and the effect of Nb loading fractions. We find that inhibitory Nbs partially clamp and stiffen D1D2, generate lattice strain in the direction parallel to several Sap-Sap contacts, and promote cooperative early-stage defect formation that nucleates from D2D2 interface dissociation. These results provide a mechanistic link between local Nb-induced conformational correlations and lattice-scale Sap depolymerization.

## Methods

### All-Atom Molecular Dynamics

Atomic structures of each isolated domain pair (D1D2, D2D3, D3D4, D4D5, D5D6) were extracted from the solved atomic model of monomeric Sap (PDB: 6HHU). Structures for domain-domain interfaces (D1D6, D2D2, D3D6, D5D5) were extracted from the Sap lattice structure solved by integrative modeling (PDB: 9G93) [48]. AAMD simulations were run with GROMACS 2022 [49] using the CHARMM36m force field [50] and TIP3P water [51] with a timestep of 2 fs; all bonds containing hydrogens were restrained using the LINCS algorithm. Each system was placed in a cubic box with a 2.0 nm buffer between the protein and all box edges, solvated in water with 150 mM NaCl, and energy minimized using the steepest descent algorithm. An NVT equilibration was performed for 5 ns at 300 K using the V-rescale thermostat [52] with a damping constant of 0.1 ps applied to the entire system, with all α-C atoms harmonically restrained with force constant 1000 kJ/mol/nm^2^ to allow the solvent and ions to relax. An NPT equilibration followed for 1 ns at 300 K using the same thermostat with damping constants of 0.5 ps for protein and solvent separately, and 1 bar maintained using the Berendsen barostat [53] with a damping constant of 5.0 ps, restraining all α-C atoms. Production NVT simulations of domain pairs were conducted at 300 K with a damping constant of 2.0 ps, saving protein configurations every 10 ps. Four independent replicas of 1000 ns each were collected per system, with the first 50 ns of each replica discarded as additional equilibration. Non-bonded domain-domain interfaces were simulated using the same GROMACS protocol as the domain-pair simulations, with four replicas of 50 ns each, saving coordinates, velocities, and forces every 5 ps. The AAMD trajectories for the Nb-D1D2 systems follows the same procedure described in ref. [25]. All resulting simulations were used as training data to map and parameterize the CG model.

Sap monomer simulations were conducted both in GROMACS [49] and on the Anton 2 [54] supercomputer. All GROMACS simulations followed the same procedure as described for the sequential domain pairs. Starting structures were the “monomeric Sap” structure (PDB: 6HHU) and the monomer rearranged into the “lattice ring” structure (PDB: 9G93). Missing sidechains were filled using CHARMM-GUI [55–57]. After equilibration, 1 μs of production NVT was run for the lattice structure and 1.2 μs for the monomeric structure. The end of each of these simulations was used as the starting structure for Anton 2 simulations using the Anton 2 protocols [54]. One replica of the lattice ring structure was run on Anton 2 for 10 μs until breaking of the D1D6 interface was observed. Three independent replicas for the monomeric structure were also run on Anton 2, spanning 3, 6, and 9 μs, respectively. Only in the 9 μs replica was release of the D3D6 interface observed. Over the total ∼30 μs collected to describe the Sap monomer in both lattice and monomer conformational states, the complete monomer-to-lattice transition was not observed, highlighting the necessity of our divide-and-conquer strategy.

### Coarse-Grained Model Construction

The CG model of Sap follows a divide-and-conquer strategy where we define and parameterize CG sites first as individual domains, then pairs of interacting domains, then as a whole-monomer in both monomer and lattice conformations, adding together the interactions at every level to reproduce the full conformational landscape of the six-domain Sap monomer. CG mappings and intra-domain interactions are defined from individual domains. Linker-domain interactions, inter-domain interactions, and Nb-Sap interactions are defined across domain pairs. Inter-domain angles, while only enforced on sequential domain pairs, are parameterized using whole-monomer simulations to incorporate changes to conformational fluctuations due to complete domain-to-domain connectivity.

CG mappings for each domain and linker were obtained independently using the K-Means Clustering Coarse-Graining (KMC-CG) method [46] as implemented in OpenMSCG [34], then applied to *α*-C coordinates extracted from the isolated domain pair (e.g., D1D2, D2D3, etc.) AAMD trajectories. For each domain or linker, KMC-CG was run over a range of candidate CG site counts, with 20 trials per count, and the optimal number of sites was selected at the elbow of the residual (*χ*^2^) vs. site-count curve. Domains were mapped to 10-20 CG sites, whereas linkers were mapped to 2-4 sites. Domain-pair, interface, and whole-monomer mappings were assembled by combining the relevant individual domain and linker mappings, then applied to the reference AAMD trajectories using center-of-mass mapping (via *cgmap*) to produce CG-mapped trajectories for parameterization.

Excluded volumes between all sites were defined using a soft potential:

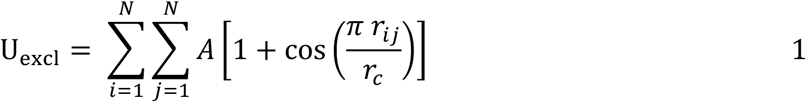

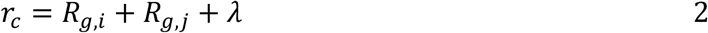

where *A* is a prefactor we set to 20 kcal/mol, *r_ij_* is the distance between sites (*i*,*j*), *R_g,i_* is the radius of gyration of all heavy atoms included in site *i*, and *λ* is a buffer distance, set to the largest it can be before soft potential energies increase in test simulations (*λ*=1.5 Å for domain-domain pairs, *λ*=0 Å for linker-interacting pairs). This buffer distance accounts for the fact that *R_g_* is prone to underestimating the effective size of each CG bead as it does not include the van der Waals radii of atoms nor possible solvent effects.

Harmonic bonds between CG sites within each domain, and between linker sites and their neighboring domains, were parameterized using heterogeneous elastic network models (hENM) [37] as implemented in OpenMSCG:

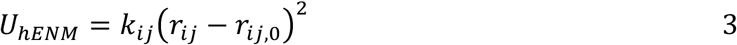

where *k_ij_* and *r_ij,0_* is the harmonic spring constant and equilibrium distance between sites (*i,j*), respectively. For each domain, an hENM was fit to the CG-mapped domain-pair trajectories over a range of cutoff distances, and the optimal cutoff was selected at the elbow of the *χ*^2^ residual vs. cutoff curve. Linker-to-domain connections were parameterized separately by applying hENM to the CG-mapped domain-pair trajectories and extracting only the bonds spanning the linker-domain boundary. No hENM bonds were placed between domains.

To capture inter-domain conformational statistics, time-lagged independent component analysis (tICA) models were trained for each sequential domain pair using CG-mapped whole-monomer trajectories. Because the thermodynamically preferred state of the Sap monomer is not known, training data were balanced to contribute equal numbers of frames from lattice-starting and monomer-starting trajectories. For each sequential domain pair (*n*, *n+1*), inter-domain angles of two types were enumerated from the CG reference structure within pair-specific distance cutoffs: angles of the form D_n_–D_n_–D_n+1_ and D_n_–D_n+1_–D_n+1_. Angles were featurized as sin/cos pairs, scaled with *RobustScaler* from *scikit-learn*, and used to train tICA models [58] with a lag time of 240 ps. The 3 highest-magnitude features from each of the top 2 eigenvectors were selected, subject to the constraint that no CG site appeared in more than one selected angle from each eigenvector. Potentials of mean force (PMFs) for each selected angle were computed via Boltzmann inversion, smoothed with a cubic spline, written to LAMMPS tabulated angle potential files [59], and enforced across all sequential domain pairs in the CG monomer (*U_angle_*).

Candidate attractive interactions at each domain-domain interface were identified using force matching [34] on CG-mapped interface AAMD trajectories. CG-mapped interface trajectories were produced by applying the relevant domain mappings to each interface system. To isolate pairwise inter-domain nonbonded forces, hENM bond forces were subtracted from the total CG forces at each frame. Specifically, LAMMPS reruns were performed on each CG-mapped trajectory with only hENM bonds active, and the resulting bond forces were subtracted from the total mapped forces, yielding nonbonded-only force trajectories. Replicas displaying clearly outlying pair distance histograms relative to the remaining replicas were excluded prior to force matching. Force matching was performed using the OpenMSCG *cgfm* utility with B-spline basis functions of order 3 and resolution 0.5 Å. Pair interaction ranges were determined per-pair from a histogram analysis: the most conservative lower bound and minimum upper bound across replicas were taken as r_min_ and r_max_, subject to a global cutoff of 30 Å. Force-matched potentials from individual replicas were combined and regularized using Bayesian ensemble regularization via *cgnes* [34]. The regularized force-matched profiles were inspected for each inter-domain CG site pair. Pairs within ∼15 Å of the interface with attractive, well-defined, monomodal force profiles were selected as candidate attractive interactions.

Attractive interactions were modeled using virtual sites (VSs), “ghost” particles that mediate anisotropic interactions between two CG sites [44]. Briefly, the VS is harmonically bound to one CG site, then allowed to interact with the partner CG site via a Gaussian potential; complete overlap of the VS and partner CG site represents the most favorable interaction between the two “real” CG sites:

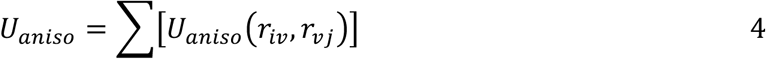

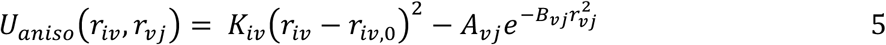

where *U_aniso_* is the (anisotropic) VS potential, *r_iv_* and *r_vj_* are the distances between CG site *i* and the VS *v*, and between the VS *v* and CG site *j*, respectively; *K_iv_* and *r_iv,0_* are the harmonic spring constant and equilibrium distance for the *i*-VS bond; and *A_vj_* and *B_vj_* are the Gaussian height and width for the VS-*j* attractive term. VS parameters were determined through a two-stage grid search. In the first stage, CGMD simulations of each isolated interface were run across a grid of *A* and *B* values for the Gaussian attractive term, and the parameter set minimizing the mean-squared error (MSE) between CG and AA-mapped interface pair distance distributions was selected as the initial estimate. In the second stage, parameters were refined in the context of a 2D periodic CG Sap lattice multiplied into a 3×3 supercell, where interfaces experience the additional mechanical load of the assembled system. A grid search over *A* values for all four interface types simultaneously was performed, again minimizing MSE to AA-mapped reference distributions. The lowest-MSE parameter set from this lattice-scale grid search was used as the final VS parameters for all CGMD simulations.

Implicit-Nb binding site correlations (*U*_3:_) were enforced as tabulated bond potentials using the same Boltzmann inversion procedure as the inter-domain angles. For each of the chosen CG site pairs, pairwise distance distributions were computed from CG-mapped (Nb-)D1D2 AAMD trajectories separately for each system. PMFs were computed with 4*π*r^2^ volume normalization, smoothed with a cubic spline with a constant restoring force applied outside the sampled region, and written to LAMMPS tabulated bond potential files. This procedure was repeated for the non-bound D1D2 system and each of the 11 Nb-D1D2 systems, yielding one bond table set per system for use as implicit-Nb potentials in the CG lattice.

The complete CG potential (**Fig. 1B**) is then:

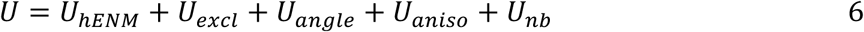

### Coarse-Grained Molecular Dynamics and Analysis

CG Sap lattices were constructed by assembling 100 monomers in a 5×10 supercell using Moltemplate [60], with each monomer comprising 102 real CG sites and 4 VSs (one per interface type), totaling 106 sites per monomer. The simulation box was triclinic with periodic boundary conditions, with initial dimensions chosen to match that of the Sap C2-symmetric dimer unit cell scaled into the supercell. Prior to depolymerization simulations, an anisotropic NP_xy_T equilibration of the non-bound lattice with in-plane *P_xx_* = *P_yy_* = 0 bar was performed to obtain stress-free box dimensions, ensuring that any strain observed in subsequent simulations could be attributed to Nb loading rather than residual lattice strain from the reference structure.

All CGMD simulations were run in LAMMPS [59] with a timestep of 50 fs. During equilibration, VS potentials were applied with elevated *A* constants (40, 40, 60, and 40 kcal/mol for D1D6, D2D2, D3D6, and D5D5 interfaces, respectively) to maintain interface integrity regardless of instantaneous monomer geometry. Final VS parameters were applied after equilibration was complete.

Each system was equilibrated in three stages: (1) energy minimization with force and energy tolerances of 10^-4^ kcal/mol/Å and 10^-6^ kcal/mol respectively; (2) lattice compression from the initial supercell dimensions to the pre-determined stress-free box dimensions over 30,000 steps in the NVT ensemble with a Langevin thermostat (300 K, damping time 10 ps); and (3) further relaxation for 1,000,000 steps in the NVT ensemble under the same thermostat. Production simulations were run for 4x10^8^ timesteps in the NVT ensemble at 300 K, with configurations saved every 50,000 steps. Three independent replicas were collected per system. Defect counts were averaged over the final 5x10^7^ timesteps of each production run.

For the bond stiffness analysis, we define the equilibrium bond distance, *r_0_*, as the expectation value of *r* across the probability distribution of each bond for that system, and quantify stiffness as the inverse of the *r*-distribution variance:

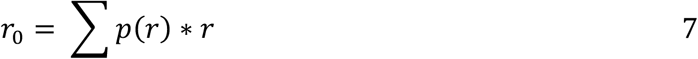

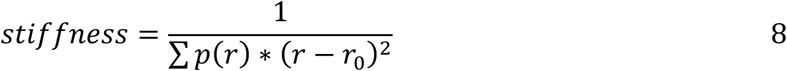

where *p*(*r*) is the probability of the bond length at distance *r*.

For the lattice strain analysis, each 100% Nb-loaded system was independently energy minimized and equilibrated under anisotropic NP_xy_T conditions (*P_XX_* = *P_YY_* = 0 bar, *τ_P_* = 100 ps, *T* = 300 K) for 100,000 steps. To prevent out-of-plane drift, which would introduce artifacts into the measured Δ*x* and Δ*y* values, a subset of CG sites (CG sites 20, 44, 54, 78, 90) was tethered to the *xy*-plane using harmonic wall potentials (*K* = 50 kcal/mol/nm^2^ between *z* = 552-558 *Å*) in the *z* direction. Equilibrium box dimensions were recorded at the end of each NPT run and compared to the stress-free reference dimensions.

### Machine learning and SHAP analysis

The hybrid classifier-regressor feed-forward neural network (FFNN) was implemented in TensorFlow [61] to identify inter-domain pair distances most predictive of Nb-induced Sap lattice depolymerization. Input features consisted of all 225 pairwise distances between the 15 D1 and 15 D2 CG sites computed from CG-mapped (Nb-)D1D2 AAMD trajectories and scaled using *RobustScaler* from *scikit-learn* [34, 62]. Frames were pooled across all replicas and randomly shuffled at the frame level, then split 60/20/20 into training, validation, and test sets, with 71,992, 23,996, and 23,996 samples, respectively.

The combined training loss was:

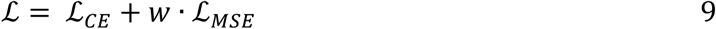

where ℒ_CE._ is the categorical cross-entropy on the classifier output, ℒ*_MSE_* is the mean squared error between the predicted probability of inhibition (p_inh_) and the probability-mapped experimental inhibitory activities at 8 *μ*M for each inhibitory Nb (mapped to [0.5,1.0], with non-inhibitory and non-bound systems set to 0), and *w* = 3 is the regression loss weight. The regression loss weight was chosen to be at the elbow of MSE between target and predicted p_inh_ over weight.

The model architecture consisted of an input layer (225 units), 1-3 hidden layers with ReLU activations, a 3-class softmax output layer (non-bound, non-inhibitory, inhibitory), and a Lambda layer extracting the inhibitory class probability p_inh_ as a regression output. Architecture hyperparameters (number of hidden layers, units per layer from [32, 64, 128, 256]) were selected via Bayesian optimization [63] with early stopping (patience = 30, maximum 100 epochs per trial). The final architecture consisted of 2 dense layers, each with 256 units. The best architecture was retrained at a reduced learning rate (10^-5^, Adam optimizer [64]) for up to 1000 epochs with early stopping (patience = 100) and the model saved when validation loss decreased. **Table S1** reports the test statistics. Final validation loss was 0.112, with a test classification accuracy of 1.00.

Per-feature SHAP importances [65] were computed using DeepExplainer on the classifier output, with 1,000 background samples drawn from the training set. SHAP values were computed over 10,000 samples and the analysis was repeated 5 times independently.

### Selection of implicit-nanobody inter-domain correlations

Ranking of candidate implicit-Nb “bonds” (i.e., pairwise distance correlations between CG sites in D1 and D2) was conducted using three methods: standard SHAP, correlation-grouped SHAP, and a Jensen-Shannon divergence (JSD)-maximizing analysis. Naively, scores for the standard SHAP bond set are calculated using the mean absolute SHAP values for the inhibitory node, averaged across 5 runs. The bond set was sorted (i.e., ranked) in descending order based on scores. The extracted bond set (for CGMD simulation) is the set of top 5 bonds, after which there is a “spectral gap,” i.e., a notable reduction in scores (see **Fig. 2B**).

**Figure 2.**
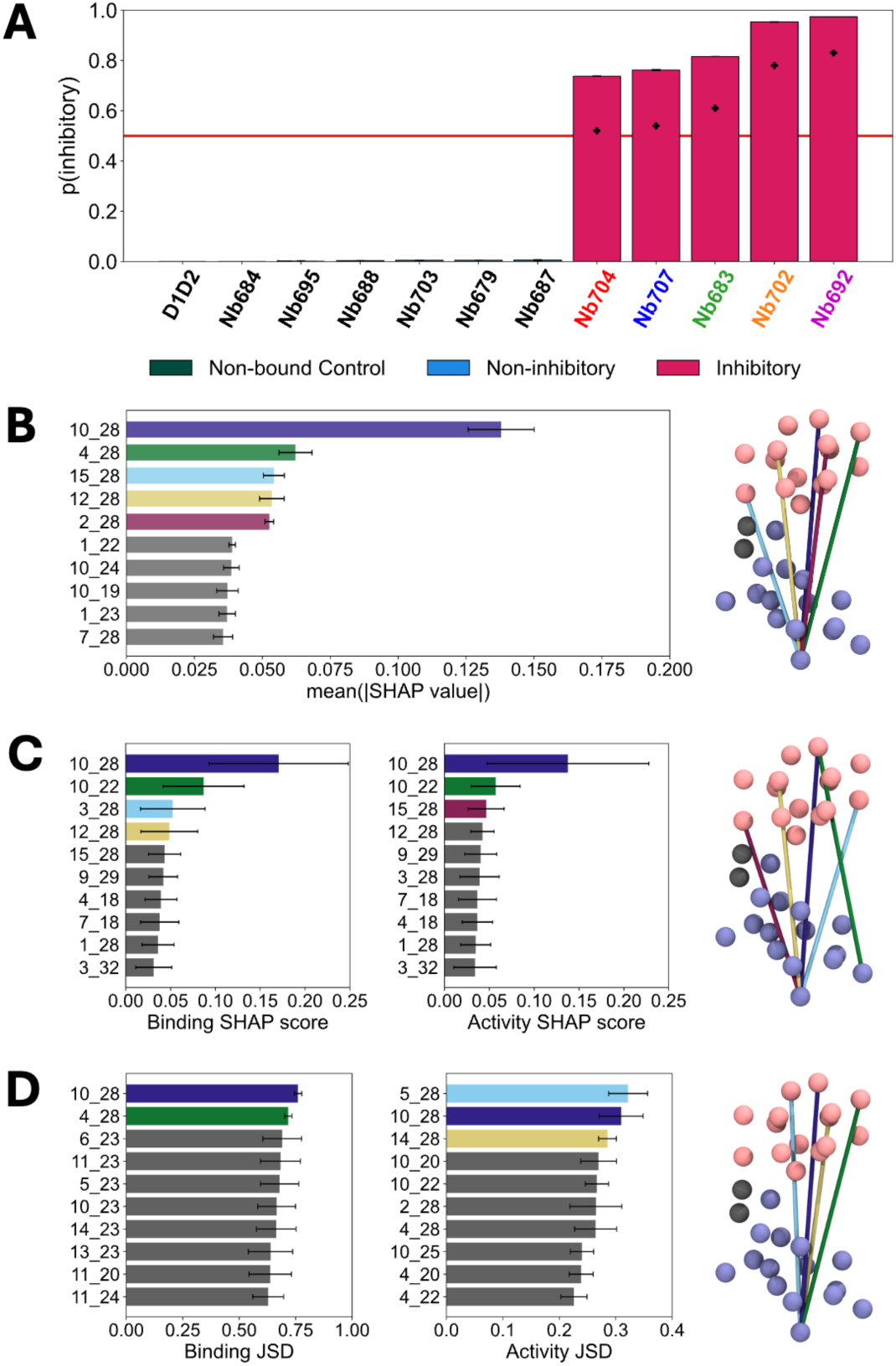
A) FFNN-predicted probability that the bound Nb (x-axis) is inhibitory. The non-bound probability is shown in green, non-inhibitory Nbs are shown in blue, and inhibitory Nbs are shown in red. Black crosses indicate the regressor target probability-mapped inhibitory activities. B) Top 10 SHAP scores (left), with the top 5 bonds rendered onto the binding site (right). Error bars indicate SEM across 5-fold cross-validation SHAP analyses. CG sites are shown using space-filling beads, where D1 is pink, D2 i ice-blue, and the linker is dark grey. Plotted bar colors match the associated bond colors. C) Top 10 Grouped SHAP scores for binding (left) and activity (center) behaviors, with the representative bonds from the top 5 groups shown on D1 and D2 (right). Error bars indicate SEM across all bonds included in each correlated group. D) Top 10 JSD scores for binding (left) and activity (center), with the top 4 pairs shown on D1 and D2 (right). Error bars indicate SEM across simulation replicas.

For the correlation-grouped SHAP analysis, we first identified correlated bonds by computing Pearson correlations between all pairs of bonds followed by grouping via a complete-linkage graph of bonds based on absolute Pearson correlations greater than 0.75. Grouped-SHAP scores were then calculated for binding and activity scores separately:

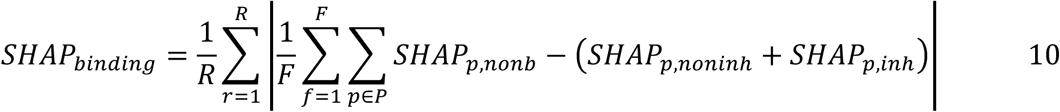

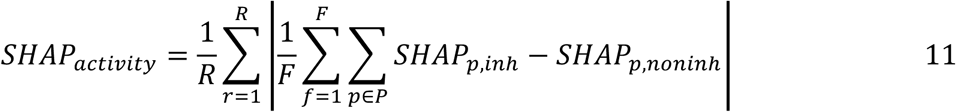

where *R* is the number of SHAP replicas, *F* is the number of frames, *P* is the set of bonds within that correlated group, and *SHAP_p,nonb_*, *SHAP_p_noninh_*, and *SHAP_p_inh_* are the frame-wise SHAP values for bond *p* for the non-bound, non-inhibitory, and inhibitory classification nodes, respectively. The bond set from this ranking is the set of representative bonds from the union of high-scoring groups in the binding (top 4) and activity (top 3) rankings, respectively, before their spectral gaps (see **Fig. 2C**). Representative bonds for each group were chosen as the bond with the highest sum SHAP_binding_ and SHAP_activity_ score within that group.

Finally, the JSD-maximizing bonds are the bonds with the largest JSD between the bound and non-bound (“binding,” top 2) or inhibitory and non-inhibitory (“activity,” top 3) datasets, averaged over each MD replica (see **Fig. 2D**). The JSD measures the similarity between two probability distributions, with JSD=0 indicating perfect overlap. Therefore, maximizing JSD selects bonds with distributions that deviate the most between bound vs non-bound or inhibitory vs non-inhibitory states.

To convert the conformational statistics of each extracted bond to bond potentials for use during CGMD, we applied Boltzmann inversion to the pairwise distance probability distributions for each bond in each (Nb-)D1D2 system, then enforced the resulting potentials as tabulated bonds in the CG model. These bonded potentials provide the “implicit Nb” contribution to the CG model. PMFs for all bonds are shown in the Supplemental Information (**Figs. S1-S3**).

## Results

**Fig. 1** summarizes the multiscale strategy used to test whether Nb-associated D1D2 correlations are sufficient to drive Sap lattice disassembly. Briefly, we developed a hybrid classifier-regressor ML model trained on CG-mapped AAMD simulations of the D1D2 binding site to identify the inter-domain (between D1 and D2) pair distances most predictive of binding and inhibitory activity (**Fig. 1A**). Candidate pairs identified either by SHAP, correlation-grouped SHAP, or JSD analysis (see *Methods*) were converted into tabulated bond potentials and enforced in a bottom-up CG model of the full Sap monomer (**Fig. 1B**). These monomers were assembled into large 2D periodic lattices (**Fig. 1C**), where the carried potential changes when Sap is unoccupied and when considered bound to each of the investigated Nbs. Therefore, when the potentials are trained from non-bound D1D2 simulations, they represent non-bound Sap; when trained on each of the Nb-D1D2 datasets, they implicitly represent each of the Nbs (hereafter referred to as an “implicit Nb”). We then apply these implicit Nbs at varying fractions to simulate the depolymerization assay (**Fig. 1C**).

### Data-driven analyses identify candidate D1D2 correlations associated with Nb inhibitory activity

Because we did not know *a priori* which specific conformational motions drive Nb-induced depolymerization, we leveraged ML to connect conformational motions to their labeled phenotypes. We developed a hybrid classifier-regressor ML model that labeled binding site conformational states according to the bound character of each system (**Fig. 1A**). Pair distances from Nb-bound D1D2 AAMD simulations were mapped to the CG resolution and classified into non-bound, non-inhibitory-bound, and inhibitory-bound states. In parallel, the inhibitory probability was biased towards the experimental inhibitory activities of each sample’s Nb (linearly mapped to the probability of inhibition or p_inh_=[0.5-1]). This hybrid objective was designed to preserve the distinction between the three classes while weakly retaining the relative activity differences within the inhibitory class. **Fig. 2A** shows the predicted p_inh_ for all trajectories of each (Nb-)D1D2 system. The ML model predicts near-zero p_inh_ for the non-bound (“D1D2”) and all non-inhibitory systems. All inhibitory systems (in red) have p_inh_ > 0.5, indicating that the classifier successfully recognized them as inhibitory. The black crosses denote the probability-mapped experimental activities for each inhibitory Nb. Where the classifier would bias inhibitory predictions for Nbs to 1 or 0 (depending on the class), the observed p_inh_ predictions are reduced toward the regressor targets, indicating that the regressor node was able to preserve the qualitative activity trend between Nbs. The final ML predictions exceed the regressor targets for all inhibitory Nbs, confirming that the regressor did not dominate over the classifier, which would otherwise collapse predictions onto the six mapped activity values.

After model training, we performed a per-feature SHAP importance analysis on the ML model to isolate the binding site pairs that are most predictive of binding and inhibitory-Nb activity, shown in **Fig. 2B**. To determine how many pairs to enforce in the CG model, we examined the sorted SHAP values per pair (i.e., SHAP “rankings”) of the inhibitory node, looking for “spectral gaps” that might define the minimal bond set required to encode Nb-binding and activity. We observed three clear gaps in the SHAP rankings (after 1, 2, and 5 pairs), each defining a candidate set of increasing size. Preliminary testing revealed that the top 5 pairs (shown on the right side of **Fig. 2B**) were required, after conversion to bonds, to observe the phenotypic effects of each Nb in lattice simulations, so we used this bond set for the first of our three candidate bond sets, which we denote as the “SHAP” set.

The SHAP analysis we used for the first bond set makes the important assumption that all pair distance features are independent of each other. Due to the inter-connected nature of CG sites within the binding site, most of our pair distances are highly correlated with other nearby pairs. Indeed, 4/5 of the top SHAP pairs are highly correlated with each other (|Pearson ρ| > 0.75). How best to account for correlated datasets with hundreds of features in SHAP analyses is still an open question [66]. Here, we corrected for correlated pairs by grouping them by their Pearson correlations (see *Methods*) and averaging SHAP values across all pairs within each correlated group. Additionally, we split the scoring into their average contributions to binding classification (non-bound vs. both bound classes) and activity classification (non-inhibitory vs. inhibitory). We then selected the top 5 correlated groups observed collectively in the binding and activity classifications. From these correlated groups, we selected a representative pair from within each group, based on the joint score from both the binding and activity classifications, and denote this bond set as our “Grouped SHAP” set (**Fig. 2C**). This set differs from the first SHAP set in that each pair encodes a different non-/weakly-correlated motion, with 3/5 new motions not represented in the SHAP set.

Finally, to compare the ML-based rankings with a simpler distribution-based criterion, we generated a third bond set using a JSD analysis [67]. We calculated the JSD for each pair distance distribution between non-bound vs. bound datasets (“Binding JSD”) and non-inhibitory vs. inhibitory datasets (“Activity JSD”). We note that this does not account for differences in depolymerizing activity within the inhibitory class or consider correlations between pairs. Here, we take the top 4 pairs before errors increase and scores become indistinguishable (**Fig. 2D**). This set shares the top-scoring pair CG10-CG28 with both SHAP and Grouped SHAP sets and shares the CG4-CG28 pair with the SHAP set. Three pairs in the JSD set are in the same Pearson correlated group (CG10/4/5-CG28), and the other pair (CG14-CG28) is in the group represented by CG3-CG28 in the Grouped SHAP set.

Pair-distance statistics for each bond set were then converted into CG bond potentials through Boltzmann inversion. PMFs of the top bond pair across all three methods (CG10-CG28), and the second-scoring pair in the Grouped SHAP set (CG10-CG22) are shown in **Fig. 3A** and **Fig. 3B**, respectively (PMFs for all bonds are shown in **Figs. S1-S3** in the Supplemental Information). In general, when the binding site is non-bound, the implicit Nb bonds show wide, often multimodal profiles (>1 energy minimum). This is consistent with what we expect from non-bound dynamics, since there is no other protein bound to restrict available conformational states. Upon Nb binding, these minima tend to shift to smaller distances, with the exception of non-inhibitory Nb684, which instead samples the larger of the non-bound minima. In general, greater inhibition appears to cause greater restriction of these pairs, shown by a narrowing of PMFs. Nb704 displays the lowest inhibition in experiments and likewise displays the widest PMF profile (least restriction) of the inhibitory Nbs. However, this rule does not hold for all Nbs, as the profiles for non-inhibitory Nb679 almost exactly match those of inhibitory Nb692 and Nb702 in both pairs. In general, the visible difference between non-inhibitory and inhibitory PMFs becomes smaller as associated bond ranking scores decrease. From this analysis, we concluded that each bond set describes important Nb-induced conformational changes on the binding site and could be suitable for use as implicit-Nb bonds.

**Figure 3.**
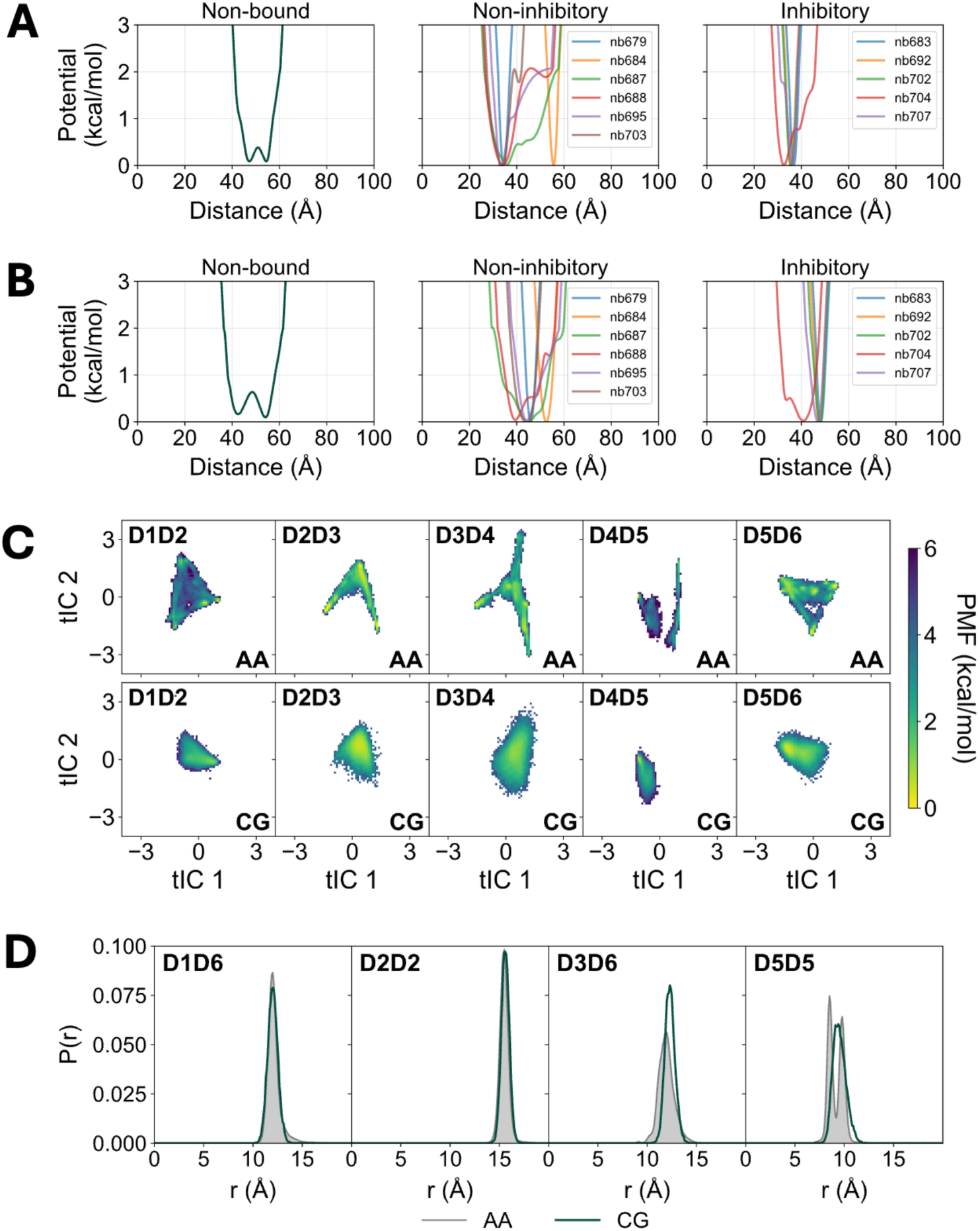
A) Potential profiles of the top-scoring implicit-Nb bond across all SHAP, Grouped SHAP, and JSD bond sets (CG10-CG28) for each Nb, separated by phenotype. B) Potential profiles of the second implicit-Nb bond in the Grouped SHAP bond set (CG10-CG22) for each Nb, separated by phenotype. C) Potential of Mean Force (PMF) profiles of sequential domain-pair angles projected into 2D-tICA space. The top row displays the training all-atom (AA) statistics, and the bottom row displays the coarse-grained (CG) statistics. Each column represents a different sequential domain pair (D1D2, D2D3, etc.). D) Comparison of CG resolution pairwise distance probability distributions describing attractive interactions at each domain-domain interface, enforced by VSs. Each interface has one VS enforcing an interaction between two CG sites. The depicted CG statistics are after optimization by grid search.

### Coarse-grained model formulation and validation

The CG model formulation for the Sap lattice follows a bottom-up coarse-graining philosophy. Each monomer consists of 6 Ig-like domains connected by 5 flexible linkers, totaling 593 residues and 9044 atoms. Due to the large conformational space likely sampled by each monomer, we used a divide-and-conquer approach. We individually simulated each sequential domain pair (e.g., D1D2, D2D3, etc.) and domain-domain lattice interface (i.e., D1D6, D2D2, D3D6, and D5D5) in AAMD, then aggregated the statistics involving each region of the protein for mapping and parameterization. This divide-and-conquer strategy allows local atomistic statistics from tractable subsystems to be combined into a lattice-scale model that retains domain rigidity, interdomain flexibility, and reversible interface binding.

Full CG model details (see **Fig. 1B**) are provided in *Methods*. Here, we briefly review the major components and verify that the CG model accurately reproduces atomistic Sap dynamics. CG-mappings were defined using KMC-CG [46] on each isolated domain or linker. Excluded volume between all CG sites was added to prevent sites from overlapping onto each other, and an hENM [37] was used to connect sites within domains and linker sites to neighboring domains. Inter-domain angles were added to reproduce the conformational motions of each sequential domain pair in the context of the whole monomer. We quantified this conformational motion using tICA models for each sequential domain pair and verified that the CG Sap monomer reproduces the major free energy minima, shown in **Fig 3C**. The CG model (bottom row of **Fig. 3C**) samples the same conformational space as the AA reference (top row of **Fig. 3C**) and shows the smoothing expected from averaging atomistic interactions to lower-resolution CG interactions.

To incorporate reversible binding of domain-domain interfaces, VSs were defined for all 4 interface types in the Sap lattice: D1D6 holding the Sap ring closed, D3D6 holding rings together along the x-axis in **Fig. 1C**, and both D2D2 and D5D5 connecting rings along the y-axis in **Fig. 1C**. **Fig 3D** shows the pairwise distance probability distributions of the reversible inter-domain interactions enforced by VSs. D1D6 and D2D2 both show exceptional overlap between the reference AA interaction (grey) and the optimized CG interaction (green). The D3D6 CG distribution is narrower and taller than the AA distribution. We attribute this to greater flexibility in the atomistic binding interface than our CG model possesses; attempts at reducing interaction strength or broadening the interaction range caused the interface to destabilize. The D5D5 AA distribution displays two peaks due to minor asymmetry at the interface. Our CG model averages this interaction in a single-peaked symmetrical interface. Together, this procedure encodes the base conformational sampling of each domain-domain junction as observed in AAMD, independent of any Nb-induced effects at the D1D2 binding site, establishing a physically grounded baseline for the monomer’s collective motions prior to Nb loading.

At this stage, the CG Sap monomer described the non-bound lattice as it included no information on the effects of bound-Nbs. Next, we incorporated the tabulated bond potentials of binding site correlations (i.e., implicit Nbs) identified by SHAP, Grouped SHAP, or JSD analyses (**Figs. 3A-B**, **Figs. S1-S3**). All monomers carried the same possible D1D2 bond terms; the non-bound or Nb-bound state is determined by the conformational statistics used to parameterize those terms. Non-bound monomers were parameterized from unbound D1D2 trajectories, while Nb-loaded monomers carried potentials derived from the corresponding Nb-bound D1D2 trajectories. Thus, Nb binding was encoded by the specific conformational changes it imposed on D1D2, instead of through explicit Nb particles or Nb-Sap contact interactions.

Finally, to obtain a periodic CG Sap lattice, we took the CG-mapped Sap dimer unit cell and replicated it into a 5x10 supercell; by default, each monomer was modeled using the non-bound D1D2 tabulated bond potentials. For a given Nb-loading fraction, we randomly selected Sap monomers in the lattice and replaced their non-bound binding site potentials with Nb-bound potentials. An example is shown in **Fig. 1C**, where blue monomers indicate that binding sites were treated with non-bound potentials, and red monomers were treated with specific Nb-bound potentials.

### Implicit Nb depolymerization assays identify phenotype-consistent binding-site correlations

To test whether the selected D1D2 correlations are sufficient to reproduce Nb action on Sap, we developed a CGMD depolymerization assay. Since the implicit Nb model enforces only a reduced set of pairwise correlations rather than a complete representation of each Nb-bound conformational ensemble, we do not expect it to quantitatively reproduce experimental activities. Instead, the assay was designed to test if selected correlations encode the dominant mechanical effect of inhibitory Nbs, such that enforcing these correlations in the Sap lattice results in the formation of inter-domain defects that we would expect prior to monomer release.

To maximize the measurable response, we constructed a periodic lattice of 100 Sap monomers at 100% implicit-Nb-loading for each Nb system and bond set, a total of 36 (12 x 3) combinations. We then ran 3 replicas of CGMD for 4x10^8^ timesteps and quantified the number of inter-domain defects averaged across the ends of each simulation; defects were counted for the four relevant Sap interfaces, i.e., D1D6, D2D2, D3D6, and D5D5. We then compared the ranked extent of defect formation with the experimental activities using Kendall’s τ_b_ correlation coefficient, since we expect the number of defects to correlate directly (although, not necessarily linearly) to inhibitory activity.

**Fig. 4** shows the average number of defects delineated for each inter-domain interface for all three bond sets per implicit Nb, including the non-bound Sap control. In general, implicit Nb loading was able to induce lattice defects above the non-bound Sap control, indicating that local binding-site correlations propagate to the scale of the assembled lattice. However, the three bond sets differed in how specifically they reproduced inhibitory vs non-inhibitory behavior. The SHAP set (**Fig. 4A**) produced a clear activity-dependent response amongst the inhibitory Nbs, with defect counts increasing monotonically with experimental activity and with an overall rank correlation of τ_b_=0.636. In addition, the non-bound control and 4/6 of the non-inhibitory Nbs remained weakly perturbed with few defects (≤ 7). However, non-inhibitory Nb679 and Nb703 displayed defect counts comparable with the most inhibitory Nbs, suggesting that this bond set missed important inhibitory correlations. This behavior is consistent with the fact that we had selected individual pairs from a correlated feature space. When mechanistically related motions are distributed across many pairwise distance fluctuations, the apparent importance of any one pair can be diluted or displaced onto partially redundant features.

**Figure 4.**
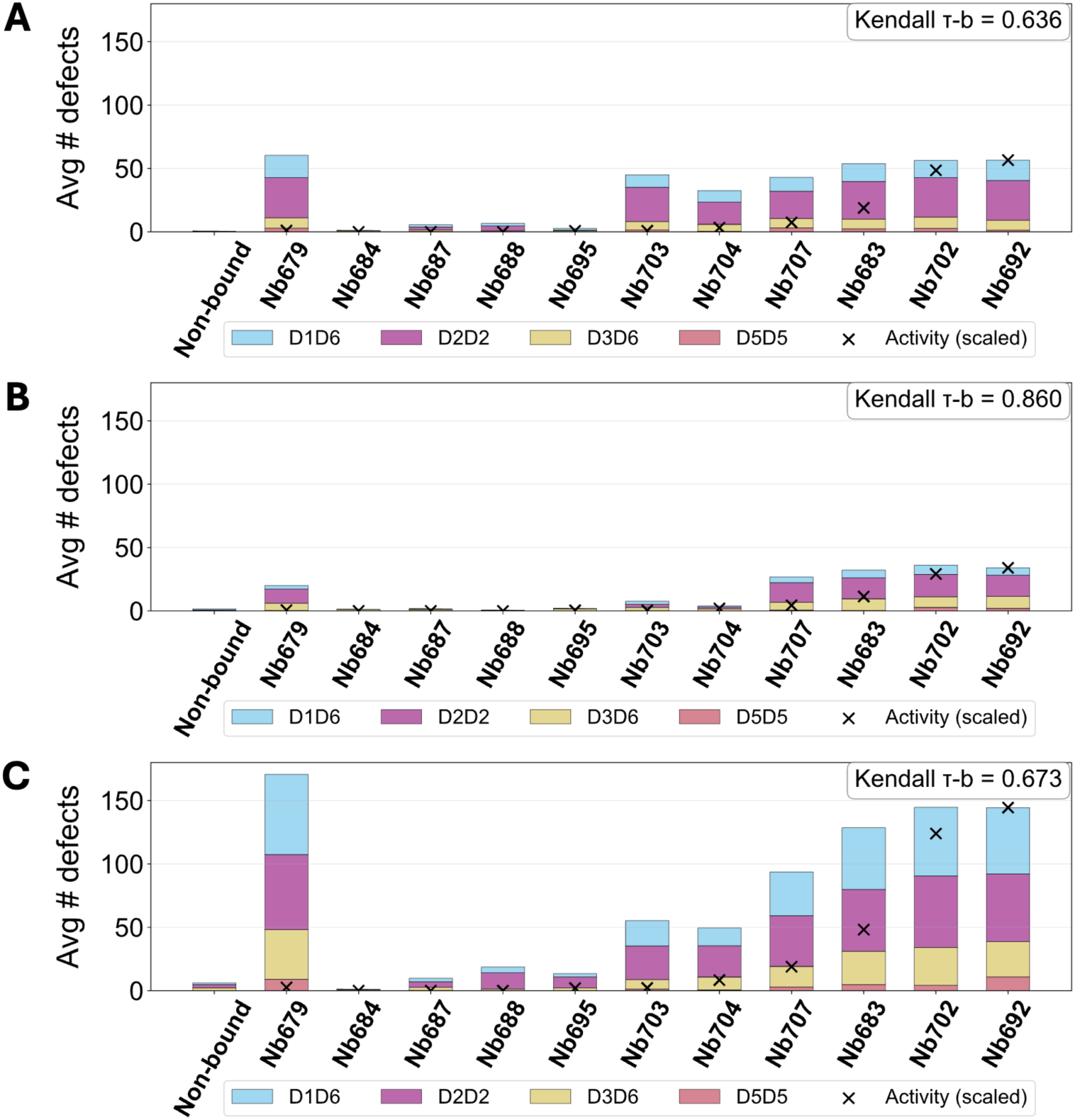
Average number of defects (y-axis) in the last 5x10^7^ CGMD timesteps for 100% loading of each Nb (x-axis), separated by interface type for A) SHAP bond set, B) Grouped-SHAP bond set, and C) JSD bond set. Crosses indicate the experimental activities linearly mapped to number of defects observed in Nb692. Kendall τ_b_ coefficients report how well the relative experimental activity ranking of Nbs is reproduced by the observed number of defects (higher is better).

Grouping correlated features after SHAP analysis improved this phenotype-level discrimination. The Grouped SHAP bond set (**Fig. 4B**) produced the fewest defects overall but showed the most consistent agreement with experimental activity rankings, yielding the highest rank correlation of τ_b_=0.860. In particular, the apparent false-positive responses of non-inhibitory Nb679 and Nb703 were reduced relative to the SHAP bond set, and the weakly inhibitory Nb704 was shifted toward the low-defect regime expected from its experimental activity. Therefore, although the Grouped SHAP set does not generate the largest absolute defect signal, it provides the most consistent reduced representation of Nb activity. We interpret this muted but more selective response as a consequence of an enforced bond set that only includes a single representative bond for correlations associated with Nb binding. Therefore, the bond set should underrepresent the full effect of inhibitory Nbs while preserving the most robust phenotype-associated motions.

Interestingly, the JSD bond set (**Fig. 4C**) showed the opposite behavior by producing the highest number of defects overall, while reproducing the expected monotonic increase for inhibitory Nbs with a rank correlation of τ_b_=0.673. However, this stronger response came at the cost of specificity. Within the non-inhibitory Nbs, there is strong defect formation on Nb679 and Nb703, and weak but non-negligible defect formation on noninhibitory Nb687, Nb688, and Nb695. These results suggest that JSD identifies structural correlations that are highly capable of destabilizing the lattice when enforced in isolation, but it also overestimates Nb activity when compensating or stabilizing correlations are absent from the implicit Nb model. In this sense, the JSD bond set appears to be overly aggressive in that it detects a depolymerization signal, but not one that is sufficiently selective for inhibitory phenotypes. We therefore did not use the JSD bond set for mechanistic interpretation of Nb action.

The behavior of both non-inhibitory Nb679 and Nb703 highlights an important limitation of the present reduced representation. Nb679 produces significant defects across all three bond sets. Given the similarity of Nb679’s implicit-Nb potentials to inhibitory Nb692 (**Figs. 3A-B**, **Figs. S1-S3**), this behavior is not surprising. Nb703 displays a similar but weaker depolymerizing trend. These results suggest that both Nbs contain local correlation signatures that resemble inhibitory Nb binding, even though they were classified experimentally as non-inhibitory. One likely possibility is that the current reduced model successfully captures destabilizing correlations at the binding site but omits compensatory correlations distributed across lower-ranked or correlated D1D2 pairs. We also cannot exclude the possibility that the experimental classification of these two Nbs is incomplete or context-dependent — a concern worth noting given that experimental labels are used directly in model training, meaning any misclassification propagates into the learned phenotype landscape.

Among the three bond selection strategies, the Grouped SHAP bond set provides the best balance between activity ranking and phenotype specificity, achieving the highest rank correlation with experimental activity (τ_b_=0.860). The bond set partially corrects the over-active responses of Nb679, Nb703, and Nb704, while preserving defect formation consistent with experimental phenotypes for 10/12 (Nb-)Sap systems. We therefore used the Grouped SHAP bond set for subsequent mechanistic analysis of Nb-action on Sap lattices, where we investigated how defects nucleate, propagate, and reorganize the lattice over time.

### Time-resolved defect analysis reveals a sequential pathway for lattice disassembly

We next inspected the development of defects over CG time using the Grouped SHAP bond set, focusing on one representative non-inhibitory and inhibitory Nb-Sap system. For non-inhibitory Nb684, defects appeared spontaneously and sporadically throughout the simulation but remained short-lived and spatially localized, as seen in **Fig. 5A**. Thus, while non-inhibitory implicit Nbs appear to transiently perturb Sap interfaces, these fluctuations do not accumulate into a propagating lattice disassembly mode. Indeed, as seen in **Fig. 5B**, the lattice is in a fully intact state at the end of the simulation.

**Figure 5.**
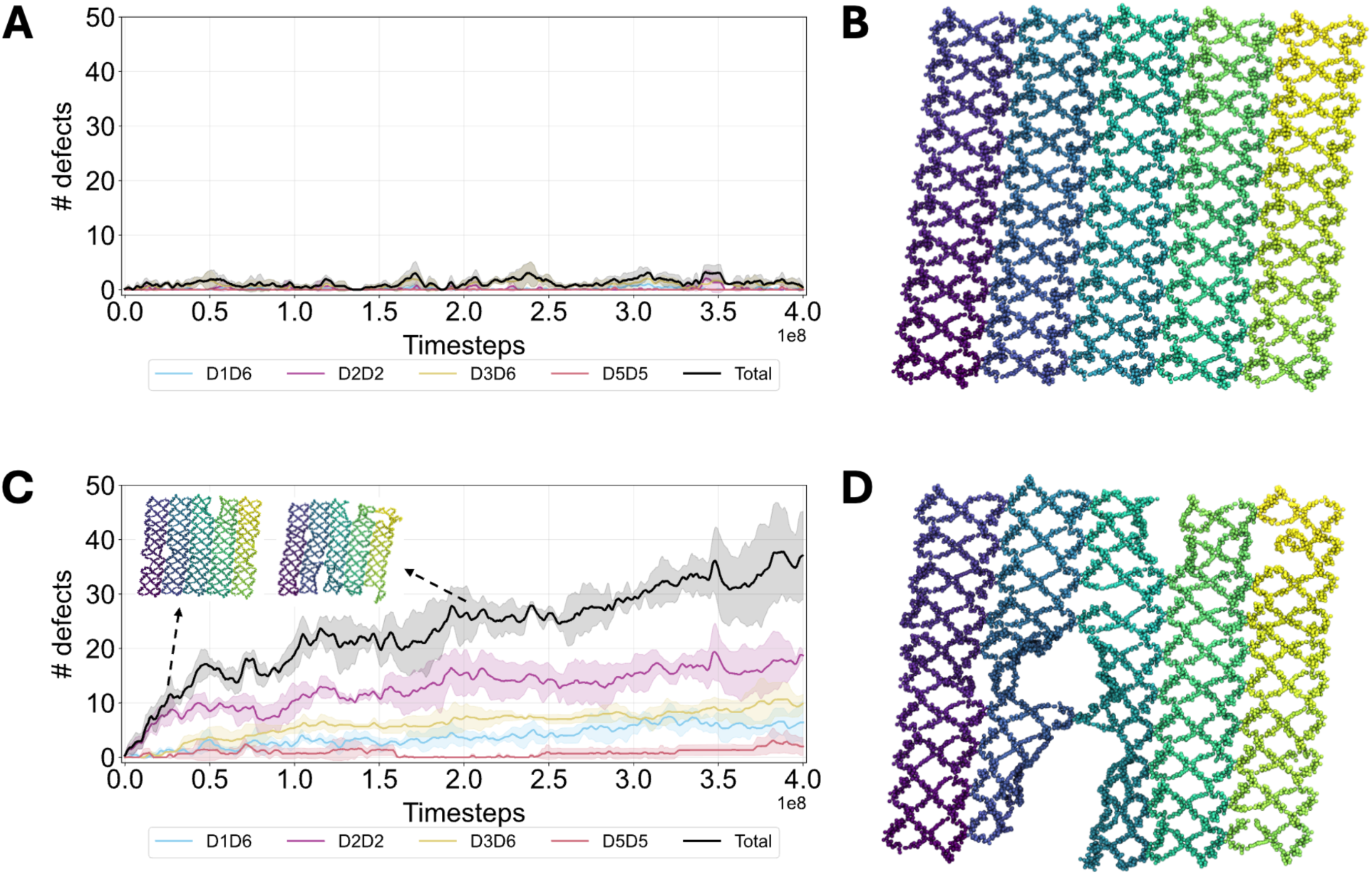
A) Time-series plot of the number of defects for 100%-loaded non-inhibitory Nb684 over CGMD timesteps. The shaded region indicates standard error of the mean (SEM) across 3 independent replicas. B) Snapshot of the final frame of 100%-loaded Nb684. Each ring is a single CG Sap monomer, colored by atom index. C) Time-series plot of the number of defects for 100%-loaded inhibitory Nb692 over CGMD timesteps. D) Snapshot of the final frame of 100%-loaded Nb692.

The inhibitory Nb692 system showed a qualitatively different trajectory (**Fig. 5C**). Defects appeared first at the D2D2 interface (a contact aligned to the y-axis in **Fig. 1C)** and grew rapidly until ∼0.25x10^8^ timesteps. Only after this initial D2D2 failure did defects begin to accumulate at the D1D6 (ring closure interface) and D3D6 (aligned to y-axis) interfaces. By the midpoint of the simulation, D2D2 contacts scattered around the lattice had opened, forming small seams. By the end of the simulation (shown in **Fig. 5D**), the D2D2 seams had widened further and were held together only by a narrow set of D1D6 and D5D5 contacts. The majority of D1D6 (ring opening) defects can be found along the other scattered seams, suggesting that the nearby D2D2 contact stabilizes the Sap ring conformation.

The spatial organization of these observed defects suggests a sequential mechanism for Nb-induced depolymerization. The earliest and most extensive lattice failure occurs at the D2-D2 interface, indicating that inhibitory Nb-induced rigidification of the D1D2 binding site first destabilizes the interface between neighboring Sap rings. As the D2D2 contacts dissociate, strain propagates to adjacent D2D2 interfaces above and below the initial defect, causing a D2D2 seam to extend throughout the lattice. Once this seam exposes an edge, nearby D1D6 and D3D6 contacts begin to fail, consistent with opening of the Sap ring and dissociation from adjacent Sap monomers. The final contact to destabilize would be D5D5, which is furthest away from the Nb binding site and likely a result of the fluctuations gained throughout the other Sap domains upon their release from binding interfaces.

Although full monomer dissociation was not observed on the present simulation timescale, the observed defect sequence is consistent with an early-stage depolymerization pathway. We therefore interpret the Grouped SHAP implicit-Nb simulations as capturing the initiation and propagation of lattice failure. Longer simulations (at least an order of magnitude longer) would likely be required to observe complete dissociation from the exposed lattice edges.

### Local D1D2 clamping and stiffening by nanobodies converts into lattice-scale strain

While observing the rigidification of inhibitory-predictive pairs from the Grouped SHAP analysis, and the resultant propagation of defects in the depolymerization assay, we hypothesized that the Nb-induced rigidification of the D1D2 binding site enacts a measurable and collective mechanical strain on the lattice. First, we quantified the bond shortening (i.e., if Δ*r_0_* < 0) and stiffening effect of all implicit-Nb bonds compared to non-bound (**Fig. 6A**). Along the Δ*r_0_* axis, we observed a wide range (∼20 *Å*) of altered bond lengths with the mean bond length reduced by ∼3-5 Å, showing that Nbs cause most bonds (within the bond set) to shorten while lengthening a select few. This is consistent with the shifts in energy minima found in **Figs. 3A-B** and **Fig. S2**: the top bond (CG10-CG28) shortens as much as 17 Å, while the others vary between -6 ≤ Δ*r_0_* < 5 Å. In the case of Nb684, its increase in mean Δ*r_0_* is reflective of the Nb constraining D1D2 into the larger of its two observed energy minima when non-bound. For all other systems, the overall effect of Nb-binding is a partial clamping of D1D2. More informative is the Δ stiffness axis, where there is clear delineation between Nbs that cause significant depolymerization in our simulations and Nbs with little to no effect. Inhibitory Nb692, Nb683, Nb702, and non-inhibitory Nb679 all present stiffened bonds (0.75 < Δ stiffness < 1.5) and each open considerable defects (see **Fig. 4B**). Inhibitory Nb707 opens fewer defects and displays half the stiffness of the other depolymerizing Nbs, while the non-depolymerizing Nbs all have bonds with little to no stiffening. From this analysis, we observe that depolymerization only occurs when the binding site is both stiffened and partially clamped. We do note that the stiffening effect is non-monotonic with inhibitory activity, which could be attributed to our earlier observation that our implicit-Nb CG model is a reduced, incomplete description of Nb action.

**Figure 6.**
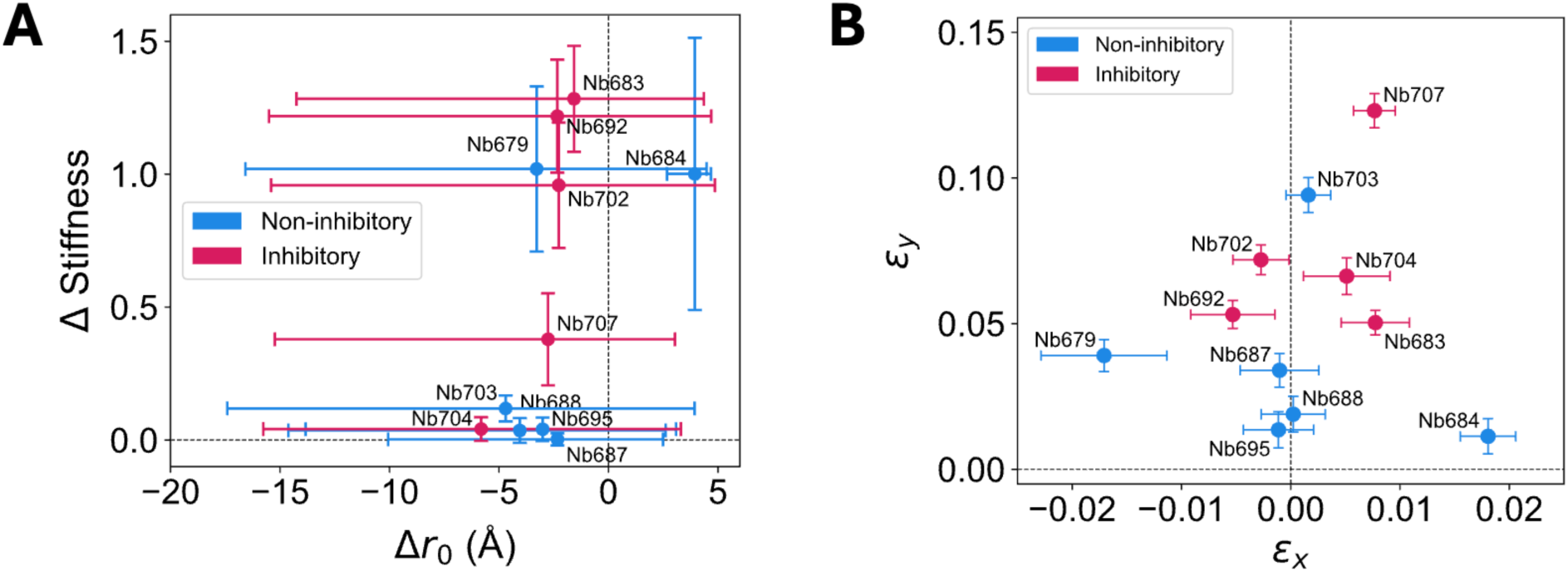
A) Change in mean bond stiffness (Δ Stiffness) over change in mean effective bond length (Δ*r_0_*) for all Nb bonds in the Grouped SHAP bond set with reference to that of the non-bound bound. Error bars indicate the min/max of Δ Stiffness and Δ*r_0_* across all bonds within each bond set. B) Strain in the x- (*ε_x_*) and y-dimensions (*ε_y_*), indicating the change in equilibrium *x* and *y* box dimensions at *P* = 0 atm divided by the non-bound *x* and *y* box lengths for the listed Nbs. Each lattice is under 100% loading of each (Nb)-D1D2 from the Grouped SHAP bond set. Error bars indicate the SEM of strain across the final 5 equilibrium strain measurements.

Next, we asked whether local D1D2 clamping and stiffening imposes a preferred direction for collective strain on the Sap lattice. We ran anisotropic NP_xy_T equilibrations on each of the Sap lattices at 100% loading, setting the target in-plane pressure *P_xx_ = P_yy_* = 0 bar. The resulting change in equilibrium box dimensions indirectly reports the direction and magnitude of the intrinsic stress introduced by each implicit Nb; an increase (decrease) in any direction implies a tensile (compressive) stress along that direction. We therefore measured the change in relaxed *x*- and *y*-dimensions for each Nb-loaded system relative to that of the non-bound reference.

**Fig. 6B** shows the strain in the *x*- and *y*-dimensions, which is the change in equilibrium *x*- and *y*-dimensions divided by the non-bound equilibrium box lengths, for each implicit Nb, colored by experimental phenotype. The stiffening and partial clamping of the binding site by Nbs (see **Fig. 6A**) translated into small effects on the *x*-dimension, straining it within ±0.01. Instead, the Nb effect was far more notable on the *y*-dimension with all Nbs causing expansion. However, according to this analysis, the extent of *y*-expansion does not seem to quantitatively correlate to experimental inhibitory activity (or observed number of defects). It seems that all Nbs with observed defect formation (Nb679, 707, 683, 702, and 692) exhibited a critical amount of expansive y-strain *ε_y_* >0.04. This *y*-directed stress is consistent with the sequence of defect formation observed in **Fig. 5**, where D2D2 contacts fail first, followed by D1D6 and D3D6 defects near exposed seams; all three of these contacts dissociate in the same direction as this intrinsic stress. In other words, the NPT relaxation assay identifies the direction in which implicit Nb loading introduces expansive mechanical strain, which is subsequently relieved through dissociation of interfaces oriented in the same direction.

### Nb-induced defect formation exhibits positive cooperativity

The depolymerization assay, bond stiffness, and lattice strain analyses together suggest that the rigidification induced by inhibitory Nbs, while local to the binding site, propagates into a larger-scale strain that promotes D2D2 defects near and around the Nb-bound Sap. This mechanism suggests possible cooperativity between bound Nbs, where increasing the number of bound Nbs would increase the number of local strain sources, such that defects become more likely to nucleate in mechanically weakened regions of the lattice. To test this hypothesis, we quantified the degree of cooperativity for each Nb by fitting the number of defects across Nb loading fraction to the Hill equation:

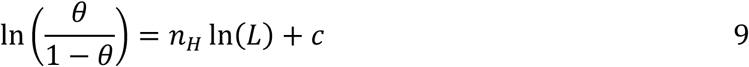

where θ is the fraction of defects, n_H_ is the Hill coefficient, *L* is the Nb loading fraction, and *c* is a fit constant. Intuitively, 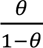 represents the odds that an interface will dissociate.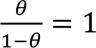 or In 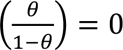 then represents the crossover point where defective and intact interfaces are equally likely. When n_H_ > 1, Nb-induced formation of defects is positively cooperative, meaning that with each Nb added, defects form more easily. Conversely, when n_H_ < 1, defect formation is negatively cooperative with respect to Nb binding, indicating that binding more Nbs decreases the likelihood of forming defects.

We repeated the depolymerization assay for every Nb with a fractional loading from 10% to 100%. **Fig. 7A** and **Fig. 7B** show the Hill equation fits for non-inhibitory and inhibitory Nbs, respectively, and **Fig. 7C** summarizes the fit *n_H_* values. All Nbs that exhibited substantial depolymerization under 100% loading (see **Fig. 4**) also displayed weak positive cooperativity (2 > *n_H_* > 1). This result is consistent with our hypothesized defect-propagation mechanism: adding more Nbs nucleates more D2D2 defects, raises the probability that neighboring defects interact, and allows local disruptions to coalesce into extended seams. Non-inhibitory Nb679 again follows inhibitory-like behavior with positive cooperativity, which is consistent with its inhibitory-like implicit-Nb bonds (**Fig. 3** and **Fig. 6**); as discussed earlier, we suspect this behavior reflects a limitation of the current model with reduced implicit-Nb representation. By contrast, the fit *n_H_* < 1 values for the non-depolymerizing Nbs should not be interpreted as true negative cooperativity. The defect likelihoods are estimated from very low (< 5) defect counts across all loadings, within error of each other, making them unreliable estimates. The meaningful distinction is therefore between systems that generate persistent defects and those that remain effectively intact. Only the depolymerizing Nbs show reproducible positive cooperativity, supporting a model in which increasing Nb occupancy amplifies early-stage defect formation, likely by introducing multiple local sources of lattice stress. However, since full monomer dissociation was not observed within the timescale of our simulations (see **Fig. 5**), our predicted *n_H_* values only reflect cooperativity in the early stages of depolymerization (and likely, a lower bound) rather than the complete cooperative response associated with full lattice disassembly.

**Figure 7.**
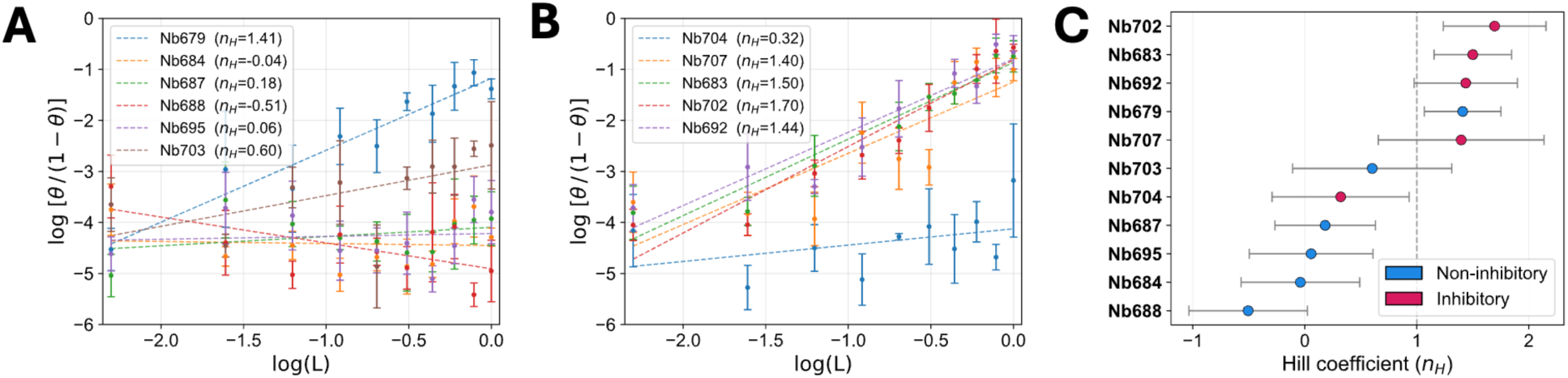
A) Hill plot of all non-inhibitory Nbs, showing the log-odds of defect formation as the log of the loading fraction *L* changes. Error bars indicate the SEM over 3 independent replicas. Dotted lines indicate the linear fit of the log-log trend, and the fit slope (*n_H_*) is shown in the legend. B) Hill plot of all inhibitory Nbs. C) Hill coefficients (*n_H_*) for each Nb. Error bars indicate the 95% confidence interval of the log-log fit.

## Discussion

Nbs are promising but still emerging tools for antivirulence therapeutics [68]. Many Nbs act through direct functional blockade, for example by neutralizing toxins, blocking virulence factors, preventing pathogen-host binding, or conformationally locking a target protein into an inactive state [68–72]. These mechanisms are well suited to design strategies that optimize affinity, specificity, or multivalency. However, some inhibitory mechanisms depend less on whether a binder occupies a particular site and more on how binding reshapes the conformational ensemble of the target. This distinction is especially relevant for large protein assemblies, where local binding can produce effects that are only expressed at the scale of the assembled structure.

The Sap-depolymerizing Nbs studied here fall into this latter category because they alter the internal mechanics of an assembled protein lattice. They bind locally to the D1D2 hinge, yet the experimental phenotype is lattice-wide depolymerization. Thus, the relevant mechanism of action cannot be inferred from binding pose or affinity alone. In this study, we sought to determine whether microscopic correlations learned through AAMD and ML are sufficient to reproduce the macroscopic effects of Nbs on assembled Sap lattices. We found that the direct atomic details of the Nb-Sap contact are not strictly required as explicit interactions in the lattice-scale CG model to reproduce most Nb-dependent phenotypes. Instead, local Nb binding appeared to reshape the conformational ensemble of D1D2, and a reduced representation of the altered ensemble was sufficient to transmit strain throughout the assembled lattice.

Our findings build directly on our prior AAMD/ML study [25], which showed that D1D2 fluctuations were predictive of whether a bound Nb was inhibitory. However, prediction did not establish whether those fluctuations were part of the depolymerization mechanism or merely correlated with it. Here, we tested that connection directly by converting inhibitory-predictive D1D2 correlations into implicit-Nb bond potentials and enforcing them in assembled CG Sap lattices. The selected pairs learned by the ML model are long-range correlations from opposite ends of D1 and D2 (**Fig. 2**), representing downstream signatures of atomic-level interactions at the binding site. These CGMD simulations reproduced experimental phenotypes for 10 out of 12 Nb-Sap systems, indicating that these propagated, collective correlations are sufficient to initiate phenotype-consistent lattice defects. Thus, the CGMD simulations converted the ML-identified correlations from predictive markers into testable mechanical perturbations that drive lattice-scale remodeling.

The observed depolymerization mechanism is inherently mechanical. Sap monomers form a C2-symmetric dimer unit cell, with D2D2 and D5D5 binding along the short *y*-axis, D3D6 binding along the long *x*-axis, and D1D6 holding monomeric rings closed (as oriented in **Fig. 1C**). In the Grouped SHAP simulations, inhibitory Nb692 caused rapid, persistent opening of D2D2 interfaces, whereas non-inhibitory Nb684 produced only transient, self-healing interface openings. These D2D2 openings are parallel to the *y*-axis stress identified by the NPT relaxation assay (**Fig. 6B**). Once D2D2 opens and an edge is exposed, Sap rings appear to lose the lateral stabilization normally provided by D2D2 contacts, making D1D6 ring closure less favorable. Upon ring opening, D3D6 loses its geometric support and destabilizes as well. Together, these observations suggest a sequential lattice failure pathway where Nb-induced D1D2 rigidification introduces tensile stress, initially leading to dissociation of D2D2 contacts, followed by exposed seams that promote loss of D1D6 and D3D6 contacts. Our simulations most directly resolved the initiation and propagation of lattice failure, yet complete monomer release was not observed. Instead, based on our observations, we expect that on a sufficiently long timescale the thermal fluctuations of open rings on exposed edges would eventually pull D5D5 interfaces apart, fully releasing Sap monomers into solution, consistent with the release of free Sap monomers observed experimentally.

The similarity in implicit-Nb potential profiles in **Figs. 3A-B** and the increase in stiffness with inhibitory activity observed in **Fig. 6A** suggest that all of the Nbs induce similar conformational changes to D1D2, but only when enforced strongly enough does depolymerization proceed. The strain analysis clarifies this point. The Nb systems that depolymerize in CGMD are those that both stiffen the D1D2 bond set and generate larger *y*-directed stress in the lattice. In the NPT relaxation assay, non-depolymerizing systems occupy a lower-strain regime, whereas defect-forming systems span 0.04 < *ε_y_* ≤ 0.12 (**Fig. 6B**). We interpret this separation as evidence that inhibitory-like Nbs impose a critical amount of stress along the direction relieved by loss of D2D2, D1D6, and D3D6 contacts.

In the cooperativity analysis (**Fig. 7**), we found that all Nbs that cause lattice depolymerization in our simulations also display weak positive cooperativity, meaning that inhibitory Nbs compound their activities with increased loading. This finding is consistent with our hypothesis that Nbs enact local strain on the Sap lattice: as the number of Nb-loaded monomers increases, the lattice contains more local sources of mechanical stress and a greater probability of forming persistent D2D2 defects. While cooperativity and the magnitude of depolymerization at saturating Nb concentrations (which we call “inhibitory activity” here) are distinct quantities, they are related. More precisely, increasing the positive cooperativity for a Nb lowers the loading fraction required to achieve a given depolymerization response (or defect formation), which is one possible strategy to increase inhibitory activity. However, we note that the Hill coefficients from the present study should be interpreted as reporting cooperativity during early-stage defect formation, as we did not observe complete lattice disassembly on our simulated timescales.

Two non-inhibitory Nbs (Nb679, Nb703) consistently produced inhibitory-like behavior during implicit-Nb CGMD simulations. These systems expose an important limitation of the reduced implicit-Nb representation. The bond sets capture dominant D1D2 correlations associated with inhibitory activity, but they may omit compensatory correlations that stabilize the full Nb-bound lattice. For highly correlated feature groups, SHAP importances can be misleading when the constituent features are only important collectively. In such cases, SHAP importance no longer reliably reflects a feature’s true contribution to the underlying phenomenon. This motivated our exploration of grouping SHAP values by correlated feature groups or taking an information-theoretic approach (e.g., JSD). Bond sets that were chosen using the SHAP or JSD analyses only represented two non-correlated motions, and both bond sets overestimated the inhibitory activity of non-inhibitory Nbs (**Figs. 4A,C**). When we collected bonds into correlated motions and re-scored according to the mean SHAP contribution of each whole group, the inhibitory activity was suppressed for Nb679 and largely eliminated for Nb703 (**Fig. 4B**), simultaneously bringing inhibitory Nbs more in line with their relative experimental activities. Therefore, grouping SHAP importances according to correlated motions provided a reduced representation of Nb action that limited the overemphasis on redundant destabilizing motions. This, however, came at the cost of total strength of Nb action on the lattice: by only enforcing a single bond for each motion, we found that total defect counts decreased. While our models are more faithful to the relative activities, they likely underestimate the total strength of inhibitory Nbs on Sap.

The clinical motivation behind this study was to investigate the mechanism by which Nbs depolymerize the Sap S-layer protein and consequently disarm *B. anthracis*. Such mechanistic insight will aid the rational design of more inhibitory Nbs. By taking a physics-based route through the conformational-activity landscape, we were able to isolate the collective motions that drive Nb-induced Sap disassembly and propose a lattice-scale depolymerization mechanism connecting local binding events to macroscopic phenotypes. The resulting ML model evaluates inhibitory activity from D1D2 conformational ensembles rather than Nb sequence directly, meaning it could, in theory, score any candidate Nb for which an AAMD simulation of the D1D2 binding site can be obtained. This type of sequence-agnostic scoring function would be suitable as a fitness function in a genetic algorithm for Nb development, as we will explore in future work. More broadly, the divide-and-conquer CG strategy combined with ML-guided parameterization is not specific to Sap: it could be applied to any large protein assembly where AAMD is tractable at the domain level but not at the assembly scale, provided a measurable macroscopic phenotype exists to supervise the ML model. Given the role of S-layers in the virulence of many pathogens, including *Clostridium difficile* and *Bacillus cereus* [13, 73], this framework could be extended to identify and exploit analogous rigidification mechanisms in other S-layer systems.

The broader opportunity is to use conformational ensemble response, rather than binding affinity or sequence alone, as a design variable for antivirulence nanobodies. This is particularly important for S-layer systems where sequence and structural specificity may require organism-specific Nbs even when a common mechanism of action (such as S-layer depolymerization) is shared. Seen from this broader perspective, Sap-depolymerizing Nbs belong to an emerging class of dynamical or assembly-disrupting antibody mechanisms, e.g., those that destabilize existing protein-protein interfaces [74, 75]. The multiscale strategy developed here provides a way to study such dynamical interventions, thereby opening a path toward antivirulence therapeutics in which nanobodies are designed to bind to virulence factors and impose collective conformational responses that ultimately disable them.

## Supplemental Materials

Additional data showing tabulated bond potentials and FFNN test statistics can be found online.

## Acknowledgements

This work was partially supported by the National Institutes of Health through grants R21AI168838 and R35GM157192. This work used high-performance computing resources from: (i) Wendian made available by the Research Computing Group at Colorado School of Mines; (ii) Anvil at the Rosen Center for Advanced Computing through allocation BIO220015 from the Advanced Cyberinfrastructure Coordination Ecosystem: Services & Support (ACCESS) program, which is supported by National Science Foundation grants #2138259, #2138286, #2138307, #2137603, and #2138296; and (iii) Anton 2 provided by the Pittsburgh Supercomputing Center (PSC) through Grant R01GM116961 from the National Institutes of Health. The Anton 2 machine at PSC was made available by D.E. Shaw Research.

## Data Availability

The data underlying this study, including simulation files, pre- and post-processing scripts, processed data, and analysis scripts, are openly available at: https://gitlab.com/pak-group/sap_depoly.

## References

1. WHO, Global antibiotic resistance surveillance report 2025. 2025.

2. Walesch, S., et al., Fighting antibiotic resistance-strategies and (pre)clinical developments to find new antibacterials. EMBO Rep, 2023. 24(1): p. e56033.

3. Tello, A., B. Austin, and T.C. Telfer, Selective Pressure of Antibiotic Pollution on Bacteria of Importance to Public Health. Environmental Health Perspectives, 2012. 120(8): p. 1100–1106.

4. Opal, S.M., et al., Antibiotic Selection Pressure and Macrolide Resistance in Nasopharyngeal Streptococcus pneumoniae: A Cluster-Randomized Clinical Trial. PLoS Medicine, 2010. 7(12).

5. Lakemeyer, M., et al., Thinking Outside the Box—Novel Antibacterials To Tackle the Resistance Crisis. Angewandte Chemie International Edition, 2018. 57(44): p. 14440–14475.

6. Rasko, D.A. and V. Sperandio, Anti-virulence strategies to combat bacteria-mediated disease. Nature Reviews Drug Discovery, 2010. 9(2): p. 117–128.

7. Allen, R.C., et al., Targeting virulence: can we make evolution-proof drugs? Nature Reviews Microbiology, 2014. 12(4): p. 300–308.

8. Totsika, M., Benefits and Challenges of Antivirulence Antimicrobials at the Dawn of the Post-Antibiotic Era. Drug Delivery Letters, 2016. 6(1): p. 30–37.

9. Dickey, S.W., G.Y.C. Cheung, and M. Otto, Different drugs for bad bugs: antivirulence strategies in the age of antibiotic resistance. Nature Reviews Drug Discovery, 2017. 16(7): p. 457–471.

10. Heras, B., M.J. Scanlon, and J.L. Martin, Targeting virulence not viability in the search for future antibacterials. British Journal of Clinical Pharmacology, 2015. 79(2): p. 208–215.

11. Fleitas Martínez, O., et al., Recent Advances in Anti-virulence Therapeutic Strategies With a Focus on Dismantling Bacterial Membrane Microdomains, Toxin Neutralization, Quorum-Sensing Interference and Biofilm Inhibition. Frontiers in Cellular and Infection Microbiology, 2019. 9.

12. Cegelski, L., et al., The biology and future prospects of antivirulence therapies. Nature Reviews Microbiology, 2008. 6(1): p. 17–27.

13. Fagan, R.P. and N.F. Fairweather, Biogenesis and functions of bacterial S-layers. Nat Rev Microbiol, 2014. 12(3): p. 211–22.

14. Johnston, E., et al., Punctuated and continuous structural diversity of S-layers across the prokaryotic tree of life. bioRxiv, 2024: p. 2024.05.28.596244.

15. Sun, Z., et al., Characterization of a S-layer protein from Lactobacillus crispatus K313 and the domains responsible for binding to cell wall and adherence to collagen. Applied Microbiology and Biotechnology, 2012. 97(5): p. 1941–1952.

16. Grogono-Thomas, R., et al., Roles of the Surface Layer Proteins of Campylobacter fetus subsp. fetus in Ovine Abortion. Infection and Immunity, 2000. 68(3): p. 1687–1691.

17. Brahamsha, B., An abundant cell-surface polypeptide is required for swimming by the nonflagellated marine cyanobacterium Synechococcus. Proceedings of the National Academy of Sciences, 1996. 93(13): p. 6504–6509.

18. Tijerina-Rodríguez, L., et al., Virulence Factors of Clostridioides (Clostridium) difficile Linked to Recurrent Infections. Canadian Journal of Infectious Diseases and Medical Microbiology, 2019. 2019: p. 1–7.

19. Sakakibara, J., et al., Loss of adherence ability to human gingival epithelial cells in S-layer protein-deficient mutants of Tannerella forsythensis. Microbiology, 2007. 153(3): p. 866–876.

20. Fioravanti, A., et al., The Bacillus anthracis S-layer is an exoskeleton-like structure that imparts mechanical and osmotic stabilization to the cell wall. PNAS Nexus, 2022. 1(4): p. pgac121.

21. Thompson, S.A., Campylobacter Surface-Layers (S-Layers) and Immune Evasion. Annals of Periodontology, 2002. 7(1): p. 43–53.

22. Dingle, K.E., et al., Evolutionary History of the Clostridium difficile Pathogenicity Locus. Genome Biology and Evolution, 2014. 6(1): p. 36–52.

23. Sogues, A., et al., Architecture of the Sap S-layer of Bacillus anthracis revealed by integrative structural biology. Proc Natl Acad Sci U S A, 2024. 121(51): p. e2415351121.

24. Fioravanti, A., et al., Structure of S-layer protein Sap reveals a mechanism for therapeutic intervention in anthrax. Nat Microbiol, 2019. 4(11): p. 1805–1814.

25. Cecil, A.J., et al., Molecular dynamics and machine learning stratify motion-dependent activity profiles of S-layer destabilizing nanobodies. PNAS Nexus, 2024. 3(12).

26. Frederix, P., I. Patmanidis, and S.J. Marrink, Molecular simulations of self-assembling bio-inspired supramolecular systems and their connection to experiments. Chem Soc Rev, 2018. 47(10): p. 3470–3489.

27. Whitelam, S. and R.L. Jack, The statistical mechanics of dynamic pathways to self-assembly. Annu Rev Phys Chem, 2015. 66: p. 143–63.

28. Lindorff-Larsen, K., et al., How Fast-Folding Proteins Fold. Science, 2011. 334(6055): p. 517–520.

29. Hénin, J., et al., Enhanced Sampling Methods for Molecular Dynamics Simulations [Article v1.0]. Living Journal of Computational Molecular Science, 2022. 4(1).

30. Salvalaglio, M., et al., Overcoming time scale and finite size limitations to compute nucleation rates from small scale well tempered metadynamics simulations. The Journal of Chemical Physics, 2016. 145(21).

31. Pak, A.J., et al., Cooperative multivalent receptor binding promotes exposure of the SARS-CoV-2 fusion machinery core. Nat Commun, 2022. 13(1): p. 1002.

32. Yu, A., et al., A multiscale coarse-grained model of the SARS-CoV-2 virion. Biophys J, 2021. 120(6): p. 1097–1104.

33. Janssens, A., et al., SlyB encapsulates outer membrane proteins in stress-induced lipid nanodomains. Nature, 2024. 626(7999).

34. Peng, Y.X., et al., OpenMSCG: A Software Tool for Bottom-Up Coarse-Graining. Journal of Physical Chemistry B, 2023. 127(40): p. 8537–8550.

35. Izvekov, S. and G.A. Voth, A multiscale coarse-graining method for biomolecular systems. J Phys Chem B, 2005. 109(7): p. 2469–73.

36. Noid, W.G., et al., The multiscale coarse-graining method. II. Numerical implementation for coarse-grained molecular models. J Chem Phys, 2008. 128(24): p. 244115.

37. Lyman, E., J. Pfaendtner, and G.A. Voth, Systematic multiscale parameterization of heterogeneous elastic network models of proteins. Biophys J, 2008. 95(9): p. 4183–92.

38. Pak, A.J. and G.A. Voth, Advances in coarse-grained modeling of macromolecular complexes. Curr Opin Struct Biol, 2018. 52: p. 119–126.

39. Majumder, A., P.G. Sahrmann, and G.A. Voth, Bottom-up Coarse-Grained Models of Asymmetric Membranes. J Phys Chem B, 2025. 129(40): p. 10333–10342.

40. Grime, J.M.A., et al., Coarse-grained simulation reveals key features of HIV-1 capsid self-assembly. Nature Communications, 2016. 7(1).

41. Pak, A.J., et al., Inositol Hexakisphosphate (IP6) Accelerates Immature HIV-1 Gag Protein Assembly toward Kinetically Trapped Morphologies. Journal of the American Chemical Society, 2022. 0.

42. Pak, A.J., et al., Off-Pathway Assembly: A Broad-Spectrum Mechanism of Action for Drugs That Undermine Controlled HIV-1 Viral Capsid Formation. Journal of the American Chemical Society, 2019. 141(26): p. 10214–10224.

43. Pak, A.J., et al., Systematic Coarse-Grained Lipid Force Fields with Semiexplicit Solvation via Virtual Sites. Journal of Chemical Theory and Computation, 2019. 15: p. 2087–2100 , pmid = 30702887 , publisher = American Chemical Society.

44. Christians, L.F., et al., Formalizing Coarse-Grained Representations of Anisotropic Interactions at Multimeric Protein Interfaces Using Virtual Sites. J Phys Chem B, 2024. 128(6): p. 1394–1406.

45. Jin, J., et al., Bottom-up Coarse-Graining: Principles and Perspectives. J Chem Theory Comput, 2022. 18(10): p. 5759–5791.

46. Wu, J., W. Xue, and G.A. Voth, K-Means Clustering Coarse-Graining (KMC-CG): A Next Generation Methodology for Determining Optimal Coarse-Grained Mappings of Large Biomolecules. J Chem Theory Comput, 2023. 19(23): p. 8987–8997.

47. Morris, E.S., et al., Multiconfigurational Coarse-Grained Molecular Dynamics. Journal of Chemical Theory and Computation, 2019. 15: p. 3306–3315.

48. Sogues, A., et al., CryoET structure of the in vitro grown Bacillus anthracis Sap S-layer. Protein Data Bank, 2024. 9G93.

49. Abraham, M.J., et al., GROMACS: High performance molecular simulations through multi-level parallelism from laptops to supercomputers. SoftwareX, 2015. 1-2: p. 19–25.

50. Huang, J., et al., CHARMM36m: an improved force field for folded and intrinsically disordered proteins. Nat Methods, 2017. 14(1): p. 71–73.

51. Jorgensen, W.L., et al., Comparison of simple potential functions for simulating liquid water. The Journal of Chemical Physics, 1983. 79(2): p. 926–935.

52. Giovanni, B., Z.-T. Tatyana, and P. Michele, Isothermal-isobaric molecular dynamics using stochastic velocity rescaling. Journal of Chemical Physics, 2009. 130: p. 74101.

53. Berendsen, H.J.C., et al., Molecular-Dynamics with Coupling to an External Bath. Journal of Chemical Physics, 1984. 81(8): p. 3684–3690.

54. Shaw, D.E., et al., Anton 2: Raising the Bar for Performance and Programmability in a Special-Purpose Molecular Dynamics Supercomputer. 2014. p. 41–53.

55. Park, S.J., et al., CHARMM-GUI PDB Manipulator: Various PDB Structural Modifications for Biomolecular Modeling and Simulation. Journal of Molecular Biology, 2023. 435(14).

56. Kong, L.Y., S.J. Park, and W. Im, CHARMM-GUI PDB Reader and Manipulator: Covalent Ligand Modeling and Simulation. Journal of Molecular Biology, 2024. 436(17).

57. Jo, S., et al., CHARMM-GUI: A web-based graphical user interface for CHARMM. Journal of Computational Chemistry, 2008. 29(11): p. 1859–1865.

58. Hoffmann, M., et al., Deeptime: a Python library for machine learning dynamical models from time series data. Machine Learning-Science and Technology, 2022. 3(1).

59. Thompson, A.P., et al., LAMMPS - a flexible simulation tool for particle-based materials modeling at the atomic, meso, and continuum scales. Computer Physics Communications, 2022. 271.

60. Jewett, A.I., et al., Moltemplate: A Tool for Coarse-Grained Modeling of Complex Biological Matter and Soft Condensed Matter Physics. J Mol Biol, 2021. 433(11): p. 166841.

61. Martín Abadi, et al., TensorFlow: Large-Scale Machine Learning on Heterogeneous Distributed Systems. arXiv, 2016: p. 1603.04467.

62. Pedregosa, F., et al., Scikit-learn: Machine Learning in Python. Journal of Machine Learning Research, 2011. 12: p. 2825–2830.

63. O’Malley, T., et al., KerasTuner. GitHub, 2019.

64. Diederik P. Kingma and J. Ba, Adam: A Method for Stochastic Optimization. arXiv, 2017: p. 1412.6980.

65. Lundberg, S.M., et al., From local explanations to global understanding with explainable AI for trees. Nature Machine Intelligence, 2020. 2(1): p. 56–67.

66. Salih, A.M., Explainable Artificial Intelligence and Multicollinearity : A Mini Review of Current Approaches. arXiv, 2024. 2406.11524.

67. Lin, J., Divergence measures based on the Shannon entropy. IEEE Transactions on Information Theory, 1991. 37(1): p. 145–148.

68. Jin, B.K., et al., NANOBODIES®: A Review of Diagnostic and Therapeutic Applications. Int J Mol Sci, 2023. 24(6).

69. De Greve, H. and A. Fioravanti, Single domain antibodies from camelids in the treatment of microbial infections. Front Immunol, 2024. 15: p. 1334829.

70. Jiang, W., et al., Small but Mighty: Nanobodies in the Fight Against Infectious Diseases. Biomolecules, 2025. 15(5).

71. Verma, V., N. Sinha, and A. Raja, Nanoscale warriors against viral invaders: a comprehensive review of Nanobodies as potential antiviral therapeutics. mAbs, 2025. 17(1).

72. Vercruysse, T., et al., An intrabody based on a llama single-domain antibody targeting the N-terminal alpha-helical multimerization domain of HIV-1 rev prevents viral production. J Biol Chem, 2010. 285(28): p. 21768–80.

73. Kirk, J.A., et al., New class of precision antimicrobials redefines role of Clostridium difficile S-layer in virulence and viability. Sci Transl Med, 2017. 9(406).

74. Koromyslova, A.D. and G.S. Hansman, Nanobody binding to a conserved epitope promotes norovirus particle disassembly. J Virol, 2015. 89(5): p. 2718–30.

75. Zheng, Q., et al., Viral neutralization by antibody-imposed physical disruption. Proc Natl Acad Sci U S A, 2019. 116(52): p. 26933–26940.

