## Supporting Information for "Collective fluctuations underlying nanobody inhibitory activity targeting *B. anthracis* S-layers revealed by multiscale simulations"

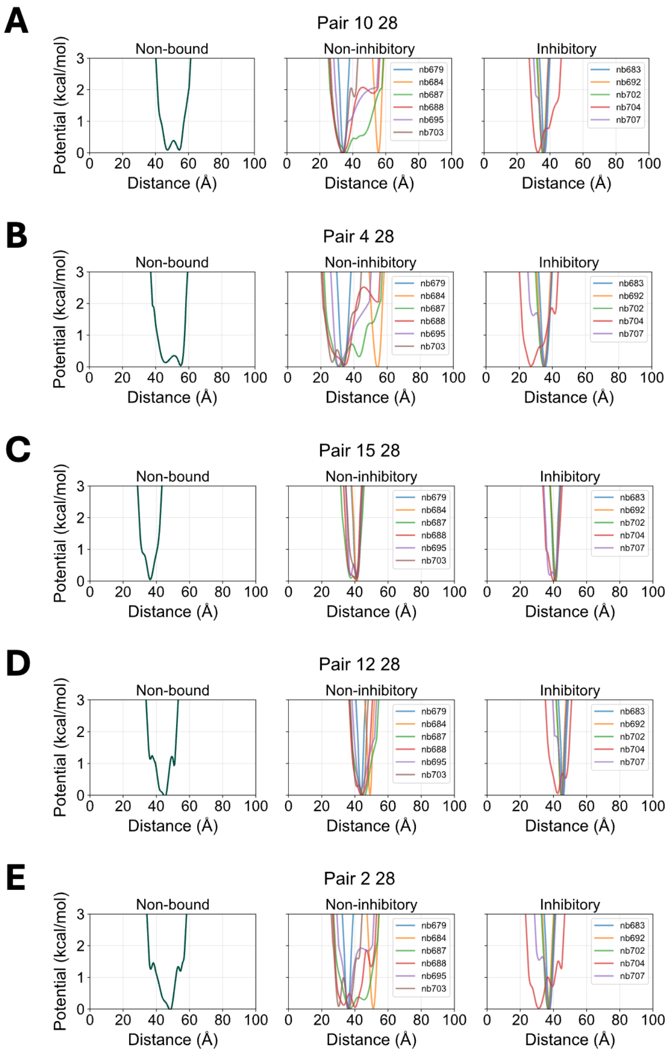


**Figure S1**. Tabulated bond potentials for the selected bonds in the SHAP bond set, separated by class. Panels (A)-(E) are ordered by decreasing importance.

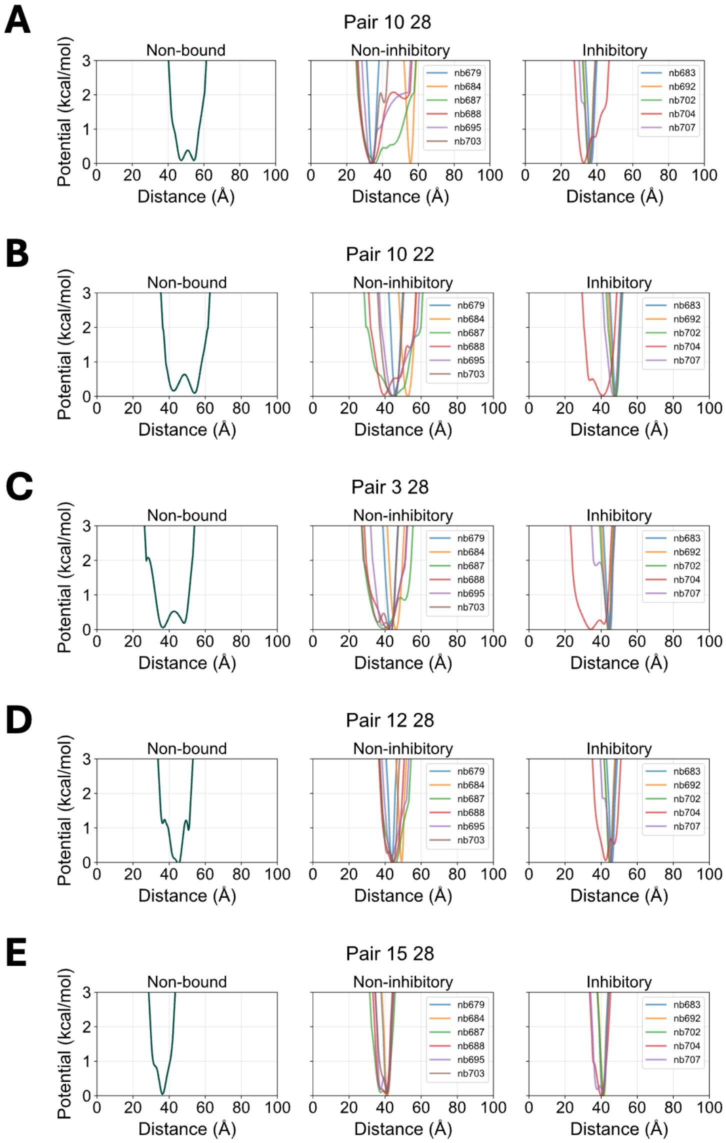


**Figure S2**. Tabulated bond potentials for the selected bonds in the Grouped SHAP bond set, separated by class. Panels (A)-(E) are ordered by decreasing importance.

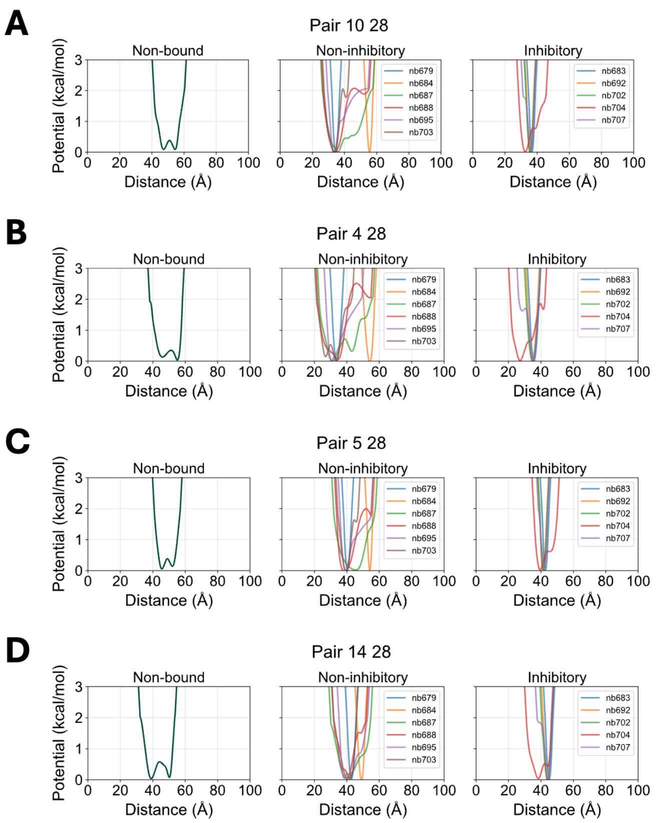


**Figure S3**. Tabulated bond potentials for the selected bonds in the JSD bond set, separated by class. Panels (A)-(E) are ordered by decreasing importance.

**Table S1**. Test accuracy statistics for FFNN hybrid regressor-classifier model.

| | **precision** | **recall** | **f1-score** | **support** |
|---|---|---|---|---|
| **nonbinding** | 1.00 | 1.00 | 1.00 | 7907 |
| **noninhibitory** | 0.96 | 0.98 | 0.97 | 8063 |
| **inhibitory** | 0.98 | 0.96 | 0.97 | 8027 |
| **accuracy** | - | - | 0.98 | 23997 |
| **macro avg** | 0.98 | 0.98 | 0.98 | 23997 |
| **weighted avg** | 0.98 | 0.98 | 0.98 | 23997 |
| **Soft classification loss (CCE)** | | | 0.0954 | |
| **Mean probability on true class** | | | 0.9305 +/- 0.1355 | |
